# A Phosphorylated Histone H2A Variant Displays Properties of Chromatin Insulator Proteins in *Drosophila*

**DOI:** 10.1101/2021.02.23.432395

**Authors:** James R. Simmons, Ran An, Bright Amankwaa, Shannon Zayac, Justin Kemp, Mariano Labrador

**Affiliations:** Department of Biochemistry and Cellular and Molecular Biology, The University of Tennessee, Knoxville, TN 37996, USA

## Abstract

Chromatin insulators are responsible for mediating long-range interactions between enhancers and promoters throughout the genome and align with the boundaries of topologically associating domains (TADs). Here, we demonstrate an interaction between proteins that associate with the *gypsy* insulator and the phosphorylated histone variant H2Av (γH2Av), a marker of DNA double strand breaks. *Gypsy* insulator components colocalize with γH2Av throughout the genome. Mutation of insulator components prevents stable H2Av phosphorylation in polytene chromatin. Phosphatase inhibition strengthens the association between insulator components and γH2Av and rescues γH2Av localization in insulator mutants. We also show that γH2Av is a component of insulator bodies, and that phosphatase activity is required for insulator body dissolution after recovery from osmotic stress. We further demonstrate a tight association between γH2Av and TAD boundaries. Together, our results indicate a novel mechanism linking insulator function with a histone H2A variant and with genome stability.

## Introduction

A highly orchestrated 3D-genome organization is necessary for the proper function and survival of eukaryotic cells. The resulting higher-order chromosome structure in this organization is driven through the establishment of long-range interactions between different regions of the chromosome. These interactions create domains that may be restricted from interacting with each other, and are based on specific interactions between chromatin-binding proteins (van Berkum *et al.*, 2010, Rao *et al.*, 2014, Fudenberg *et al.*, 2016). Chromatin insulators represent a class of protein/DNA complexes associated to specific sequences in the genome that work through two general functions: to restrict communication between enhancers and promoters through physical separation into different genomic domains and to prevent the spread of heterochromatin into euchromatic regions of the genome (Harrison et al., 1989, Geyer and Corces, 1992, Bell et al., 1999, Schoborg and Labrador, 2014, Ozdemir and Gambetta, 2019). The presence of insulators in the genome is conserved among eukaryotes, with the CTCF insulator being the only known insulator regulating the human genome (Heger and Wiehe, 2014). Insulators have recently been described to help establish the boundaries of topologically associating domains (TADs) and are often found enriched at TAD boundaries (Van Bortle et al., 2014, Fudenberg et al., 2016).

*Drosophila melanogaster* has an array of different insulator complexes, with each complex being recruited to different sequences in the genome (Bushey et al., 2009, Negre et al., 2010). Insulators were first described in fruit flies with the discovery that the scs and scs’ sequences function as boundary elements, inhibiting chromosomal position effects (Kellum and Schedl, 1991). Another insulator site, located within the *gypsy* retrotransposon, has been thoroughly characterized for its ability to regulate transcription of neighboring genes (Geyer and Corces, 1992).

A number of *gypsy* retrotransposons are present throughout the *Drosophila* genome (Hoskins et al., 2015), and insertion or transposition of *gypsy* to a new locus may interrupt local transcriptional activity and chromatin dynamics (Jack et al., 1991, Geyer and Corces, 1992). Insulator proteins are recruited to *gypsy* through a 460-bp sequence composed of 12 binding sites for Suppressor of Hairy Wing (Su(Hw)) (Geyer and Corces, 1992). The Su(Hw) protein contains an amino-terminal tandem array of a dozen C_2_H_2_ zinc finger domains (Kim et al., 1996) and a carboxy-terminal leucine zipper domain (Harrison et al., 1993) that is essential for insulator activity through interactions with other insulator proteins (Melnikova, Kostyuchenko, Molodina*, et al.*, 2018). Su(Hw) specifically recruits one of the many isoforms of Modifier of mdg4 (Mod(mdg4)67.2) through interactions between the carboxy-terminal domain of Su(Hw) and carboxy-terminal acidic domain that is unique to the *gypsy*-binding isoform of Mod(mdg4) (Ghosh et al., 2001) and through interactions between the amino-terminal end of Su(Hw) with the glutamine-rich domain of Mod(mdg4)67.2 (Melnikova, Kostyuchenko, et al., 2017). Centrosomal Protein 190 (CP190), another essential component of the *gypsy* insulator (Pai et al., 2004), was originally described through its activity during the cell cycle, dissociating from chromatin during mitotic prophase and localizing to the centrosome (Ooegema et al., 1995). In chromatin, CP190 is found as an essential part of different insulator complexes (Bushey et al., 2009) and is recruited to the *gypsy* insulator through interactions with Mod(mdg4)67.2 (Pai et al., 2004) and the amino terminal domain of HIPP1 (HP1 and insulator partner protein 1) (Melnikova et al., 2019). HIPP1 is the most recently described member of the *gypsy* insulator complex (Alekseyenko et al., 2014) and functions to stabilize the interaction between Su(Hw) and CP190, but is not required for insulator activity or transcriptional regulation (Glenn and Geyer, 2019, Melnikova et al., 2019).

Insulator activity in *gypsy* is promoted by factors such as the interchromosomal EAST protein (Golovnin *et al.*, 2015, Melnikova, Shapovalov*, et al.*, 2017) and the Chromatin-linked adapter for MSL proteins (CLAMP) (Bag *et al.*, 2019), while activity is negatively regulated by RNA-binding proteins Rumpelstiltskin (King *et al.*, 2014) and Shep (Chen *et al.*, 2019). Insulator-binding proteins in *Drosophila* can form aggregates known as insulator bodies (Gerasimova and Corces, 1998). The role of these bodies in genome organization has been debated, and functions for insulator bodies have been proposed from genome organization hubs to passive storage centers for insulator proteins (Gerasimova et al., 2000, Labrador and Corces, 2002, Golovnin et al., 2012). Previous work in our lab has demonstrated a role for insulator bodies in the cellular response to osmotic stress, with insulator proteins leaving chromatin and forming bodies in the nucleoplasm as the environment becomes more hypertonic (Schoborg et al., 2013).

Of the *gypsy* insulator proteins, Su(Hw) is perhaps the best-characterized. Mutation of *su(Hw)* is associated with female infertility (Klug et al., 1968, Harrison et al., 1993, Baxley et al., 2011, Hsu et al., 2020) and alters the cell’s response to DNA damage (Lankenau et al., 2000). Outside of the *gypsy* insulator complex, Su(Hw) binds many sites alone or in conjunction with either Mod(mdg4)67.2 or CP190 (Adryan et al., 2007, Kuhn-Parnell et al., 2008, Bushey et al., 2009, Negre et al., 2010, Soshnev et al., 2012, Baxley et al., 2017).

Apart from its role in chromatin insulator activity, Su(Hw) is also a transcriptional repressor in *Drosophila* oogenesis (Baxley et al., 2011) and regulates the expression of neuronal genes in tissues outside the nervous system, including the ovary (Soshnev et al., 2013). Mutation of *su(Hw)* leads to defects in the formation of ring canals between nurse cells, preventing the proper transport of maternal morphogens during ovary development (Hsu et al., 2015). Ovaries in *su(Hw)* mutants also exhibit malformed microtubule organizing centers and mislocalization of Gurken, a morphogen normally sequestered to one end of the oocyte to direct axis determination in embryos (Hsu et al., 2020). Similar to its role in oogenesis, Su(Hw) is critical for spermatogenesis through its activity as a transcriptional repressor in somatic cyst cells in the male germline (Duan and Geyer, 2018).

Su(Hw) also participates in the DNA damage response, possibly as part of the search for homologous sequences during homologous recombination (Lankenau et al., 2000). A role for insulators in homologous recombination-based DNA repair has been well established with mammalian CTCF, which is recruited to sites of DNA double strand breaks (DSBs) (Han et al., 2017, Hilmi et al., 2017, Lang et al., 2017). Of clinical relevance, mutation of CTCF in humans is associated with various forms of cancer (Docquier et al., 2005, Kemp et al., 2014, Canela et al., 2017, Guo et al., 2018). One of the first steps in the cellular response to DSBs is the phosphorylation of a variant of H2A known as H2AX in mammalian systems and H2Av in *Drosophila* (Rogakou et al., 1998, Baldi and Becker, 2013). H2AX is phosphorylated by ATM (Ataxia-telangiectasia-mutated) kinase and DNA-dependent protein kinase (DNA-PK) in response to ionizing radiation (Stiff et al., 2004) and by ATR (ataxia telangiectasia and Rad3-related) kinase after cells experience replication-induced genotoxic stress (Ward and Chen, 2001). Phosphorylation of H2AX leads to recruitment of numerous proteins involved in the DNA damage response (Sirbu and Cortez, 2013). Upon resolution of the DSB, H2AX is dephosphorylated primarily by PP2A (Chowdhury et al., 2005). H2Av, the sole *Drosophila* H2A variant (Baldi and Becker, 2013), combines the activities of mammalian H2AX and H2AZ. Like H2AX, H2Av is phosphorylated in response to DNA damage (Madigan et al., 2002), and like H2AZ, phosphorylation of H2Av is involved in transcriptional regulation through activation of poly(ADP-ribose) polymerase 1 (PARP1) (Kotova et al., 2011). Our laboratory previously demonstrated an accumulation of phosphorylated H2Av (γH2Av) signal in the ovaries of *su(Hw)* mutants and the presence of chromosomal aberrations in actively dividing larval neuroblasts lacking Su(Hw), suggesting a significant connection between Su(Hw) activity and genome stability (Hsu et al., 2020). Furthermore, disruption of Mei-41/ATR, a kinase responsible for phosphorylating H2Av among other targets upon DNA damage (LaRocque et al., 2007), partially rescues the defective oogenesis phenotype associated with mutation of *su(Hw)* (Hsu et al., 2020). While insulator-binding proteins have been described for their role in genome organization and regulation in *Drosophila*, the mechanisms linking their activity to DNA repair remain elusive.

In this work, we show that γH2Av is present at Su(Hw)-binding sites throughout the genome, including at *gypsy* retrotransposons and that mutation of several *gypsy* insulator components disrupts normal H2Av phosphorylation patterns. We show that γH2Av is a component of insulator bodies formed under osmotic stress and that dephosphorylation of γH2Av is required for efficient dissolution of these bodies during recovery. Chromatin immunoprecipitation (ChIP) experiments reveal extensive genome-wide colocalization between Su(Hw) and γH2Av and enrichment for both at TAD boundaries. This association also extends to insulator function as flies doubly heterozygous for *His2Av^810^* and mutant alleles of *su(Hw)* showed a partial rescue of phenotypes for *yellow^2^* and *cut^6^*, two *gypsy* insulator induced mutations. Collectively, these findings point to a model in which γH2Av works with insulators to coordinate genome function and genome-wide responses to genotoxic stress.

## Materials and Methods

### Fly stocks and husbandry

All stocks were maintained on a standard cornmeal agar fly food medium supplemented with yeast at 20°C; crosses were carried out at 25°C. The following stocks are maintained in our lab and were originally obtained from Victor Corces (Emory University): *y^2^w^1^ct^6^*; *cp190^H31-2^*/*TM6B*, *Tb^1^*, *y^2^w^1^ct^6^*; *cp190^P11^*/*TM6B*, *Tb^1^*, *y^2^w^1^ct^6^*; *w^1118^*; *su(Hw)^V^*/*TM6B*, *Tb^1^*, *y^2^w^1^ct^6^*; *mod(mdg4)^u1^*/*TM6B*, *Tb^1^*. The stock *w^1118^*; PBac(RB)*su(Hw)^e04061^*/*TM6B*, *Tb^1^* was obtained from the Bloomington *Drosophila* stock center (BDSC: 18224). These remaining stocks were provided by our lab: OR, *y^2^w^1^ct^6^*; PBac(RB)*su(Hw)^e04061^*/*TM6B*, *Tb^1^* (derived from BDSC: 18224).

### Antibodies

Rabbit polyclonal IgG antibodies against Su(Hw), Mod(mdg4)67.2, and CP190 were previously generated by our lab (Wallace et al., 2010, Schoborg et al., 2013). A rat polyclonal IgG antibody against Su(Hw) generated by our lab was also used. Antibody against the phosphorylated form of H2Av (UNC93-5.2.1) (Lake et al., 2013) was obtained from the Developmental Studies Hybridoma Bank, created by the NICHD of the NIH and maintained at The University of Iowa, Department of Biology, Iowa City, IA 52242. These antibodies were all diluted 1:1 in glycerol (Fisher Scientific, BP229-1, lot 020133) and used at a final dilution of 1:200. Secondary antibodies were all diluted 1:1 in glycerol and used at a final dilution of 1:200. The following secondary antibodies were used in this study: Alexa Fluor 594 goat anti-rabbit (Invitrogen, A-111037, lot 2079421), Alexa Fluor 488 donkey anti-rabbit (Invitrogen, A-21206, lot 1834802), Alexa Fluor 488 goat anti-guinea pig (Invitrogen, A-11073, lot 84E1-1), Texas red donkey anti-rat (Jackson ImmunoResearch Laboratories, 712-075-150), and Alexa Fluor 488 goat anti-mouse (Invitrogen, A-11001, lot 1858182).

### Immunostaining of larval tissues

Wandering third instar larvae were dissected in PBS. Tissues were immediately placed into fixative (4%, para-formaldehyde (Alfa Aesar, 43368, lot N13E011), 50% glacial acetic acid (Fisher Scientific, A38-212, lot 172788)) on a coverslip for one minute. Samples were squashed by lowering a slide on top of the sample then turning it over, placing it between sheets of blotting paper, and hitting the coverslip firmly with a small rubber mallet. Slides were dipped in liquid nitrogen, coverslips were removed, and samples were incubated in blocking solution (3% powdered nonfat milk in PBS + 0.1% IGEPAL CA-630 (Sigma-Aldrich, 18896, lot 1043)) for 10 minutes at room temperature. The slides were dried and incubated with primary antibodies overnight at 4°C in a box humidified with wet paper towels. The next day, slides were washed twice in PBS + 0.1% IGEPAL CA-630 before incubation with secondary antibodies for three hours in the dark at room temperature. Slides were washed twice in PBS + 0.1% IGEPAL CA-630, treated with DAPI solution of 0.5 μg/mL (ThermoFisher, D1306) for one minute, and washed one more time in PBS alone.

Samples were mounted with Vectashield antifade mounting medium (Vector Laboratories, H-1000, lot ZF0409) and coverslips were sealed with clear nail polish. All microscopy for immunostaining was performed on a wide-field epifluorescent microscope (DM6000 B; Leica Microsystems) equipped with a 100x/1.35 NA oil immersion objective and a charge-coupled device camera (ORCA-ER; Hamamatsu Photonics). Image acquisition was performed using SimplePCI (v6.6; Hamamatsu Photonics). Image manipulation was performed in FIJI (Schindelin et al., 2012); all contrast adjustments are linear. Images were further processed in Adobe Photoshop CS5 Extended, Version 12.0 x64. Figures were assembled in Adobe Illustrator CS5, Version 15.0.0. Statistical analyses were performed in GraphPad Prism version 8.0.0 (224) (GraphPad Software, San Diego, CA).

### Immunostaining of S2 cells

For normal control conditions, S2 cells were incubated in insect medium (HyClone SFX-Insect Cell Culture Media; Fisher Scientific, SH3027802) supplemented with penicillin (50 units/mL) and streptomycin (50 μg/mL) (Gibco, 15070063) at 25°C. To induce osmotic stress, the istonic media was replaced with hypertonic media supplemented with 250 mM NaCl (Fisher Scientific, BP358-212). Cells were treated in this hypertonic stress media for 30 minutes. Coverslips were pretreated with pure ethanol (Decon Labs, 2716) and coated with concanavalin A (Sigma-Aldrich, C5275) to help S2 cells adhere to the glass surface. Cells were pipetted onto treated coverslips and were allowed to spread and adhere for 30 minutes. After treatment, cells were fixed (4%, para-formaldehyde, 50% acetic acid) for 10 minutes at room temperature, followed by three washes with PBS buffer. Fixed cells were permeabilized with 0.2% Triton X-100 (Fisher Scientific, BP151, lot 014673) for five minutes then washed twice with PBS buffer. Cells were incubated in blocking solution (3% powdered nonfat milk in PBS + 0.1% IGEPAL CA-630) for 10 minutes at room temperature. Primary antibodies were diluted in blocking solution and samples were incubated in antibody solution overnight at 4°C in a box humidified with wet paper towels. Unbound antibodies were washed off three times with PBST buffer (0.1% Triton-X 100). Secondary antibody incubation, DAPI staining, and mounting were performed as described above.

### Okadaic acid treatment

For the isotonic control samples, S2 cells were cultured in HyClone SFX-Insect media as above and incubated in 50 nM okadaic acid (Sigma-Aldrich, O9381) for 30 minutes. The hypertonic samples were obtained by shifting S2 cells from isotonic insect media to hypertonic conditions as described above and incubating for 25 minutes. After this, the hypertonic media was supplemented with 50 nM okadaic acid and cells were incubated for five more minutes. The isotonic recovery sample was obtained by first inducing hypertonic stress for thirty minutes, including okadaic acid for the final five minutes as above, then washing out the hypertonic media twice with isotonic media containing 50 nM okadaic acid. Cells were incubated in isotonic recovery media with okadaic acid for thirty minutes. Control samples were collected throughout the process following the same protocol without addition of okadaic acid. For the polytene chromosome example, salivary glands were dissected from wandering third instar larvae and incubated in 50 nM okadaic acid for 30 minutes before fixation and squashing.

### Fluorescence intensity and colocalization analysis

Images were analyzed for the amount of each protein (i.e. the intensity of each channel) using a macro script in FIJI (Schindelin et al., 2012). The DAPI channel was used to automatically generate non-biased ROIs for each cell, which were then manually curated for extra precision. A rolling-ball background subtraction algorithm was used for all images. Intensity measurements were made using the measure function. Numerous images of polytene chromosomes were collected from each salivary gland squash. All acquisition parameters were kept constant between slides within each experiment. Colocalization was quantified using the Coloc2 plugin in FIJI. This analysis uses the Costes method (Costes et al., 2004) to determine appropriate thresholds for each channel. Results are reported in terms of Pearson’s Correlation Coefficient (PCC) (Pearson, 1895), which ranges from −1 for perfect anti-correlation to +1 for perfect correlation (Dunn et al., 2011). Another metric of colocalization is the Manders’ Colocalization Coefficients (Manders et al., 1992, Manders et al., 1993, Adler and Parmryd, 2010) for each channel – this relates how much of the signal in the green channel overlaps with signal in the red channel (M1) and how much of the signal in the red channel overlaps with signal in the green channel (M2). M1 and M2 may vary from 0, representing no overlap between signals, to 1, representing total overlap.

### ChIP-seq and TAD data analysis

The following NCBI publicly available ChIP-Seq datasets were used for the genome wide comparisons: SRX1299942 (yH2Av SRA (Li et al., 2016)), SRX046654 (Su(Hw) SRA (Chen et al., 2012)), SRX186113 (Mod(mdg4)67.2 SRA (Matzat et al., 2012)), SRX2638361 (CP190 SRA (Jox et al., 2017)), SRX2638363 (CTCF SRA (Jox et al., 2017)), and SRX511131 (HIPP1 SRA (Alekseyenko et al., 2014)). The sequencing data was uploaded to the Galaxy web platform, and the public server at usegalaxy.org was used to analyze the data (Afgan et al., 2018). Briefly, the FastQ datasets from NCBI were mapped with Bowtie2 to produce BAM files (Langmead and Salzberg, 2012). Duplicate and unmapped reads were filtered out with SAM tools. Peaks were called with Model-based analysis of ChIP-Seq (MACs) (Zhang et al., 2008). We used SeqMINER version 1.3.4 for the downstream plotting analysis (Ye et al., 2011, Zhan and Liu, 2015).

To compare the distribution of γH2Av with that of nucleosomal H2Av we used the high-resolution distribution of homotypic and heterotypic *Drosophila* H2Av nucleosomes from S2 cells obtained from a previous study (Weber et al., 2010). FastQ datasets from paired-end reads from native micrococcal nuclease–digested chromatin, enriched in homotypic H2Av (SRX019957 (Weber et al., 2010)) or heterotypic H2Av (SRX019953 (Weber et al., 2010)), were mapped with Bowtie2 to produce BAM files mapped to the dm6 *Drosophila* genome. We applied Bamcoverage to BAM files to generate H2Av Hom and H2Av Het bigwig files. SeqMINER was used to generate displays of the distribution of mean read density profiles (tags / 50 bp), using BED files to provide reference coordinates (± 10,000 bp). Bigwig files were generated by applying Bamcoverage to BAM files and peak profiles were visualized with the IGV genome browser (igv.org/app/) (Robinson et al., 2011), using *Drosophila* dm6 as the reference genome. The genomic distribution of TADs in the *Drosophila* genome (Ramirez et al., 2018) was used to produce a BED file generated with all genomic 1Kb fragments containing a TAD boundary at the center.

### Phenotypic analysis

Documentation of *y^2^* and *ct^6^* phenotypes was performed using a stereomicroscope (MZ16 FA; Leica Microsystems) equipped with a CCD color camera (DFC420; Leica Microsystems). A 150 Watt white light source (KL 1500 LCD; Leica Microsystems) set to a color temperature of 3,000 K was used for illumination. Male flies were selected soon after eclosion and aged for five days at 25°C before imaging. All images within each tissue set were collected with the same parameters using Leica Application Suite (Version 2.4.0 R1; Leica Microsystems). Abdomen images were recorded with a gamma correction of 0.5. Image analysis was performed in FIJI (Schindelin et al., 2012). Intensity of the darkest region within the fifth abdominal tergite was measured using a circular ROI with radius of 15 pixels (= 0.045 mm) (Figure 8). Intensity values from abdomens were inverted before analysis so that darker pigmentation provided a higher score. Regions of abdomens that reflected the light source were excluded from analysis. Wing areas were measured in FIJI (Schindelin et al., 2012) using the entire translucent area of the wing (except for the alula, which was sometimes lost in sample preparation) as the ROI.

## Results

### H2Av Phosphorylation Is Correlated with *gypsy* Insulator Components Genome-wide

Beyond its canonical insulator functions, the *Drosophila* insulator protein Su(Hw) has been implicated in transcriptional regulation and is required for development of testes and ovaries (Soshnev *et al.*, 2013, Duan and Geyer, 2018). Mutation of *su(Hw)* interferes with normal DNA damage repair (Lankenau et al., 2000) and lack of Su(Hw) protein results in chromosomal aberrations in developing neuroblasts (Hsu et al., 2020). While these results point to a role in maintaining genome stability, the mechanistic link between Su(Hw) and DNA repair remains uncharacterized. To investigate this possibility, we performed immunostaining of polytene chromosomes from salivary glands of third instar larvae using an antibody directed against the phosphorylated form of H2Av (γH2Av), a marker of DNA double strand breaks (Madigan *et al.*, 2002). This procedure reveals a close association between Su(Hw) and γH2Av throughout each polytene chromosome arm. Close inspection shows γH2Av in nearly all of the Su(Hw) bands (Figure 1A). Linescans of the 2R polytene chromosome show strong covariance in the fluorescent intensity between γH2Av and Su(Hw) (Figure 1A). Likewise, analysis of the immunostaining signal using the entire polytene genome shows a significant colocalization between γH2Av and Su(Hw) (Figure 1A). Quantification of colocalization of the two immunostaining signals is reported using Pearson’s Correlation Coefficient (PCC), which describes the covariance between the two signals, with positive numbers describing direct correlation between signal intensities, negative numbers representing anti-correlation of the signals, and zero representing no correlation between signals (i.e. random covariance) (Adler and Parmryd, 2010, Dunn et al., 2011). Colocalization is also reported using Manders’ overlap coefficient (MOC), which describes the amount of one signal in an image that overlaps with signal from the other channel (Manders et al., 1993). Two values are reported, one for each channel, with values ranging from 0 (representing no spatial overlap between signals) and 1 (representing complete overlap of signals).

**Figure 1.**
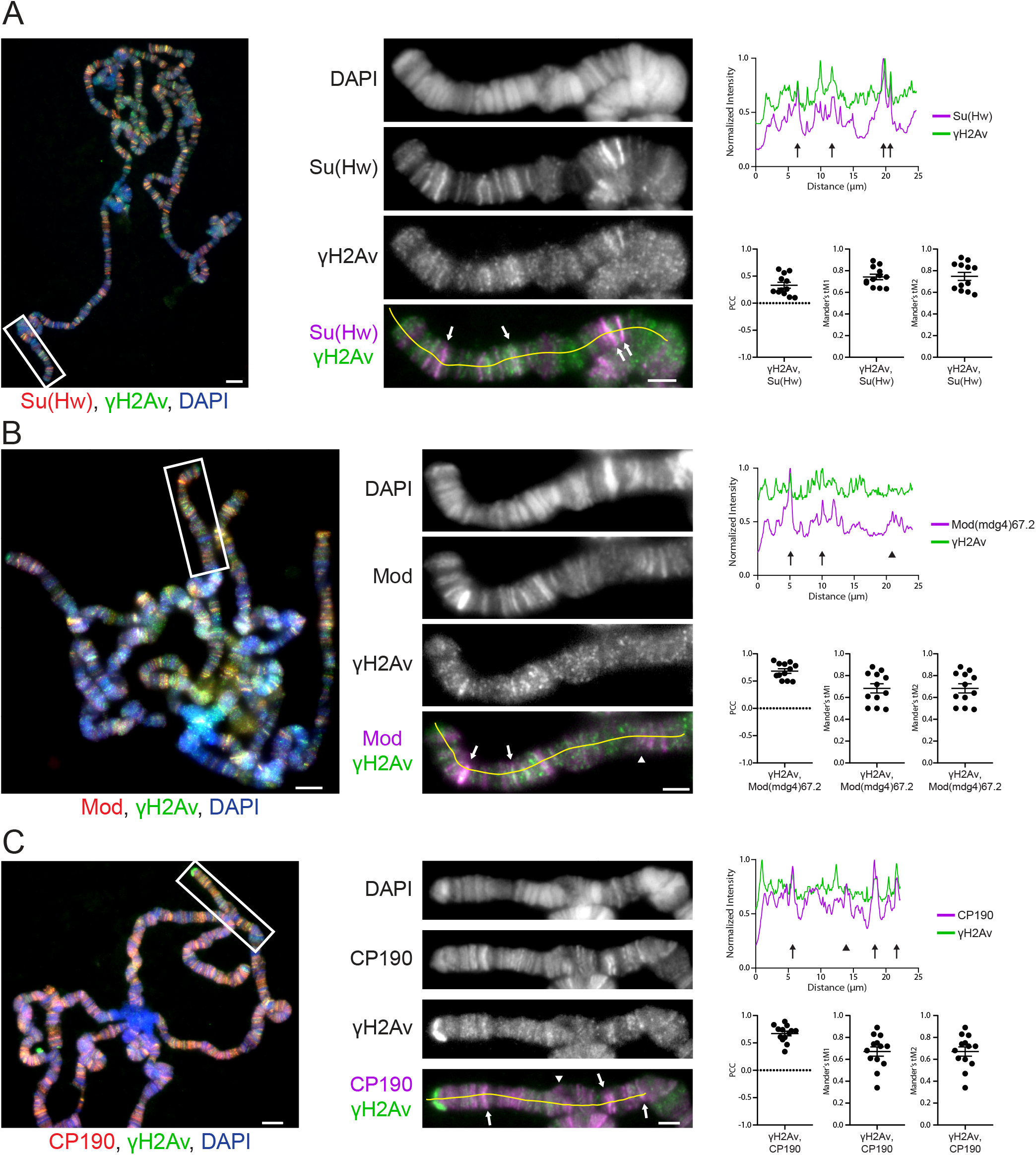
Insulator proteins colocalize with phosphorylated H2Av in *Drosophila* polytene chromosomes. **A.** Colocalization of γH2Av with Su(Hw). **B.** Colocalization of γH2Av with Mod(mdg4)67.2. **C.** Colocalization of γH2Av with CP190. Immunofluorescent micrographs of polytene chromosome squashes obtained from wandering third-instar larvae are shown on the left. Magnified insets are shown in the middle, corresponding to the white boxes in the figures on the left. Scale bars are 5 μm in the figures and 2 μm in the insets. Insets are shown as RGB merge, with DAPI on the blue channel, γH2Av on the green channel, and various insulator proteins on the red channel. Red and green channels are shown independently in grey scale and merged as magenta and green. On the right are linescans performed in FIJI (Schindelin *et al.*, 2012), corresponding to the yellow lines in the merged insets. Linescan intensities were normalized by dividing each value by the maximum intensity recorded on each channel. Arrows in the images and respective linescans denote regions of strong colocalization while arrowheads denote regions enriched for insulator proteins but not γH2Av. Pearson’s Correlation Coefficient (PCC) for γH2Av signal with each insulator protein signal is plotted, with each point representing the polytene genome of each cell. Similarly, Mander’s tM1 (the proportion of γH2Av-positive pixels that also contain signal from antibodies against insulator proteins) and tM2 (the proportion of insulator protein-positive pixels that also contain signal from the γH2Av antibody) are plotted in each cell. Error bars represent one standard error of the mean.

An extensive genome-wide association between γH2Av and Su(Hw) is illustrated by the positive PCC values and MOC values above 0.5. Given the fundamental role of γH2Av in DNA repair, the tight association of Su(Hw) with phosphorylated H2Av supports the notion that Su(Hw) is involved in maintaining genome integrity (Lankenau et al., 2000, Hsu et al., 2020). Immunostaining experiments yield similar results for Mod(mdg4)67.2, the isoform of the *mod(mdg4)* locus associated with *gypsy* insulator function (Figure 1B). Bands of Mod(mdg4)67.2 are seen to overlap with bands of γH2Av (Figure 1B) and linescans reveal a close covariance (Figure 1B). The same pattern is recapitulated when examining images and linescans of CP190 (Figure 1C), an insulator protein found at *gypsy* and CTCF insulator sites (Pai et al., 2004, Mohan et al., 2007). Quantitative analysis of signal colocalization shows strong positive correlation values when examining genome-wide signal in polytene chromosomes for both Mod(mdg4) and CP190 with γH2Av (Figure 1B, C). These findings suggest that γH2Av colocalizes with *gypsy* insulator proteins throughout the genome under normal developmental conditions.

### Phosphorylated H2Av Is Stabilized by *gypsy* Insulator Components

To further our understanding of the relationship between γH2Av and insulator complexes we asked whether or not mutation of genes coding for *gypsy* insulator complex members would affect H2Av phosphorylation. Immunostaining of polytene chromatin revealed an almost complete elimination of γH2Av in the chromatin of *su(Hw)^e04061^* mutants (Figure 2). Mutation of *cp190* also resulted in less γH2Av signal in the immunostained chromosomes, however, the reduction was not as severe as was seen in the *su(Hw)^e04061^* mutant (Figure 2). In contrast with *su(Hw)* and *cp190*, mutation of *mod(mdg4)* had no significant effect on the amount of H2Av phosphorylation in chromatin (Figure 2). These results suggest that Su(Hw) and CP190, but not Mod(mdg4)67.2, are necessary for sustaining H2Av phosphorylation and may hint at a mechanism that depends on specific interactions between H2Av with some but not all *gypsy* insulator components.

**Figure 2.**
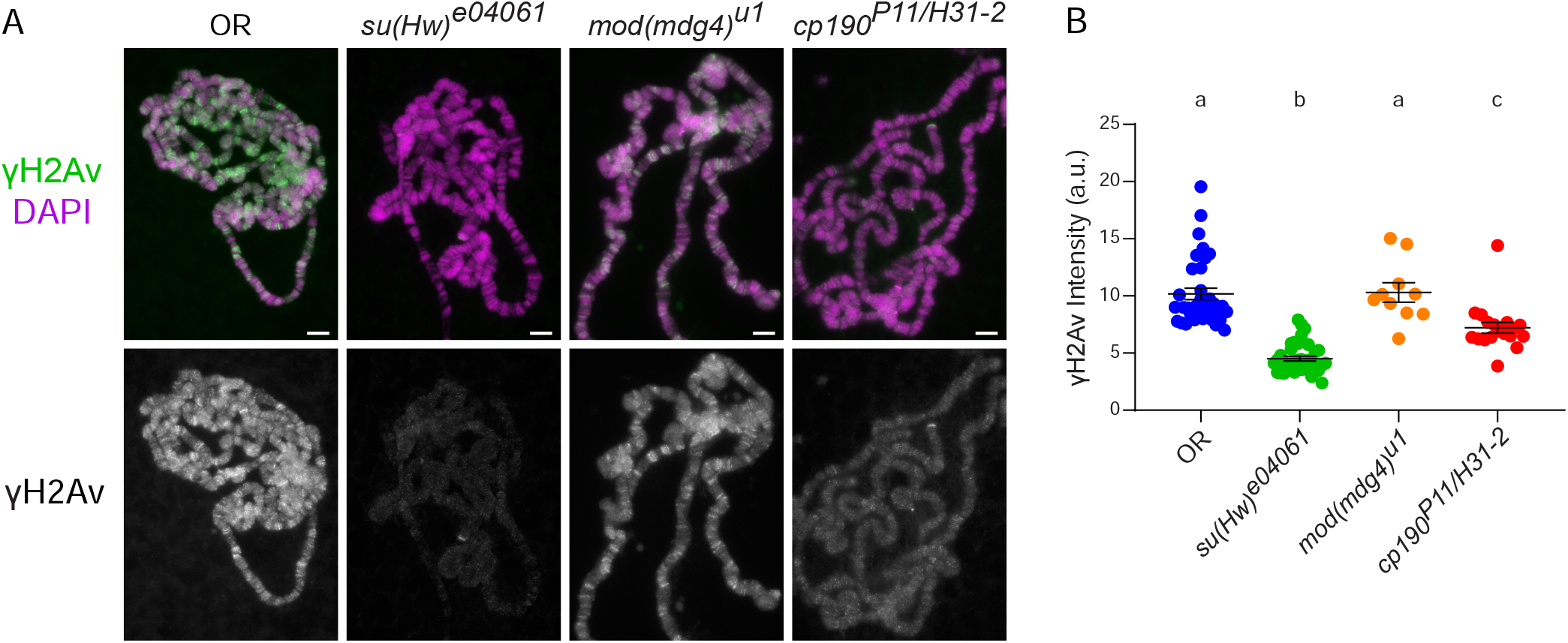
Phosphorylation of H2Av in polytene chromosomes is inhibited in insulator protein mutants. **A.** Immunostaining of polytene chromosomes from wandering third instar *Drosophila* larval salivary glands from various insulator mutant genotypes (listed above each figure). An antibody against phosphorylated H2Av (γH2Av) was used (green in the merged image, shown alone in greyscale below). Scale bars represent 5 μm. **B.** Quantification of fluorescent signals from immunostains with γH2Av shown in A. Each point represents the polytene genome of an individual cell. Error bars represent one standard error of the mean. Letiers above the data indicate statistical groupings as determined using an ANOVA performed with Games-Howell’s multiple corrections.

As interactions between insulator proteins are generally required for canonical insulator functions (Geyer and Corces, 1992, Bonchuk et al., 2015, Golovnin et al., 2016, Melnikova, Kostyuchenko, Parshikov, et al., 2018, Melnikova et al., 2019), we examined the relationship between each of these insulator proteins and γH2Av in various insulator mutant backgrounds. Notably, the colocalization between Mod(mdg4)67.2 and γH2Av was strongly reduced in the *su(Hw)^e04061^* background (Figure 3A, F). This is not surprising given both that γH2Av signal is significantly reduced in *su(Hw)^e04061^* (Figure 2) and that Mod(mdg4)67.2 does not bind at Su(Hw) sites in the absence of Su(Hw) (Ghosh et al., 2001). Linescans of polytene chromosomes and quantitative colocalization analysis show no correlation between the signals from Mod(mdg4)67.2 and γH2Av in *su(Hw)^e04061^* mutant polytene chromosomes (Figure 3A). Similarly, immunostaining for Su(Hw) in the loss of function *mod(mdg4)^u1^* mutant revealed significantly less colocalization as seen by visual inspection, linescans, and quantitative analysis (Figure 3B, F). Linescans from immunostains of the *mod(mdg4)^u1^* mutant with either Su(Hw) (Figure 3C) or CP190 (Figure 3D) along with γH2Av demonstrate a further disruption of the interaction between γH2Av and insulator complexes. The lack of colocalization between γH2Av and Su(Hw) or CP190 is reflected in decreased PCC values (Figure 3F), with negative values for CP190 indicating anticorrelation between the CP190 and γH2Av signals. Su(Hw) was also found to colocalize less with γH2Av in the null *cp190^P11-H31-2^* mutant background through the area of chromosome 2R examined (Figure 3E) and genome wide (Figure 3F). Taken together, these results indicate that stable accumulation of γH2Av in Su(Hw) insulator sites requires having the entire insulator complex intact. It remains unclear which Su(Hw) insulator protein serves as the binding partner of γH2Av, or if this colocalization requires that γH2Av interact with more than one insulator protein or other unknown protein.

**Figure 3.**
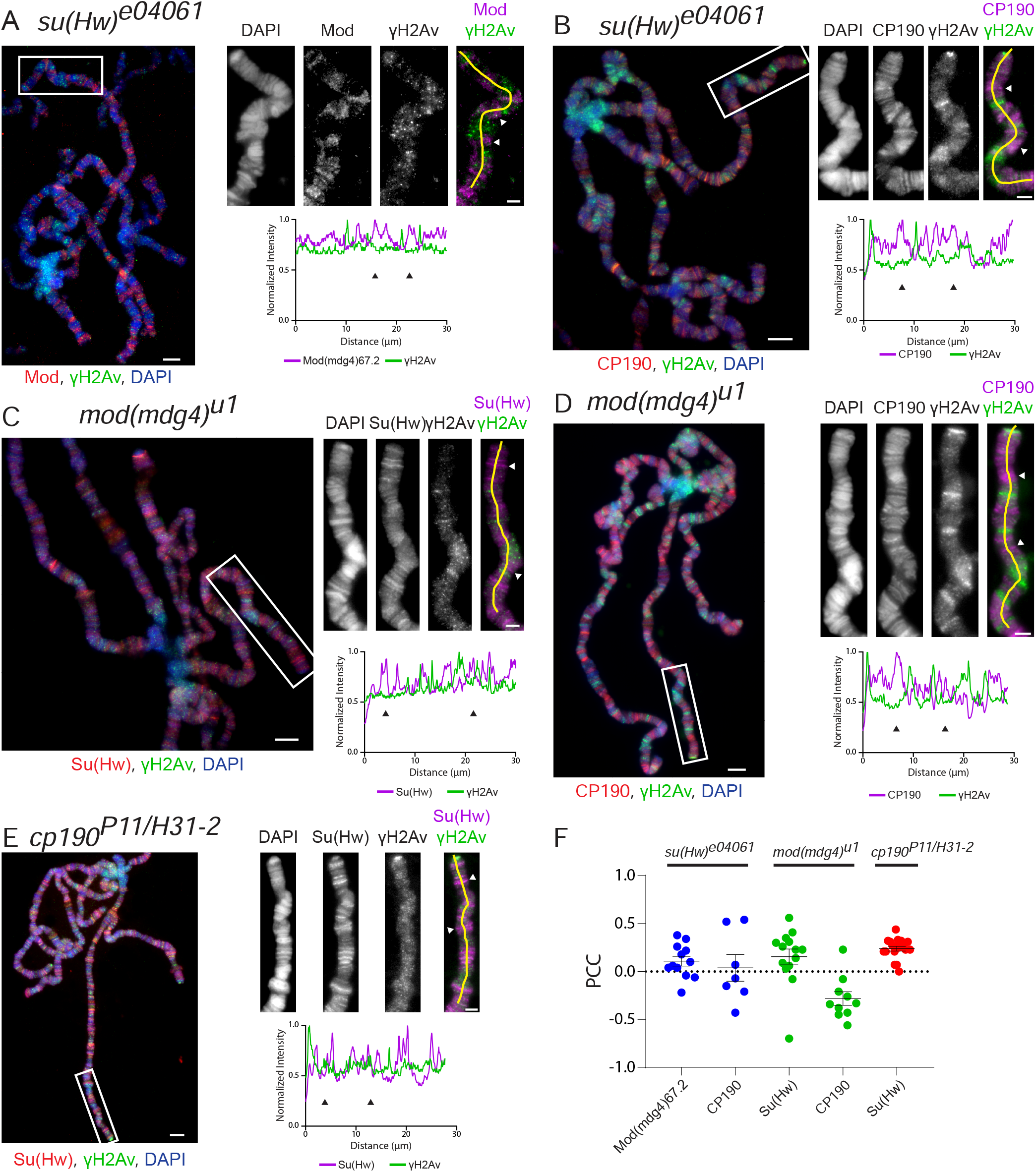
Insulator components are interdependent for their colocalization with γH2Av in *Drosophila* polytene chromosomes. **A.** Colocalization of γH2Av with Mod(mdg4)67.2 in *su(Hw)^e04061^*. **B.** Colocalization of γH2Av with CP190 in *su(Hw)^e04061^*. **C.** Colocalization of γH2Av with Su(Hw) in *mod(mdg4)^u1^*. **D.** Colocalization of γH2Av with CP190 in *mod(mdg4)^u1^*. **E.** Colocalization of γH2Av with Su(Hw) in *cp190^P11/H31-2^*. **F.** Pearson’s Correlation Coefficient (PCC) for γH2Av signal with each insulator signal is plotted, with each point representing the polytene genome of each cell. Error bars represent one standard error of the mean. PCC values are grouped by genotype (red = *su(Hw)^e04061^*, green = *mod(mdg4)^u1^*, blue = *cp190^P11/H31-2^*). Immunofluorescent micrographs of polytene chromosome squashes obtained from wandering third-instar larvae are shown on the left. Magnified insets are shown to the right of each figure, corresponding to the white boxes in the figures on the left. Scale bars are 5 μm in the figures and 2 μm in the insets. Insets are shown as RGB merge, with DAPI on the blue channel, γH2Av on the green channel, and various insulator proteins on the red channel. Red and green channels are shown independently in grey scale and merged as magenta and green. Beneath the insets are linescans performed in FIJI (Schindelin *et al.*, 2012), corresponding to the yellow lines in the merged insets. Linescan intensities were normalized by dividing each value by the maximum intensity recorded on each channel. Arrows in the images and respective linescans denote regions of strong colocalization while arrowheads denote regions enriched for insulator proteins but not γH2Av.

### H2Av Is Phosphorylated in Insulator Bodies

Upon our finding that γH2Av colocalizes with insulator proteins in the genome in an insulator-dependent manner, we next asked whether or not this interaction was maintained once insulator proteins aggregate into bodies under osmotic stress conditions. Insulator bodies represent a special case for insulator activity: proteins bound to insulator sites in the genome can leave chromatin and associate to form insulator bodies (Gerasimova and Corces, 1998). The exact purpose of these bodies remains unknown, but recent reports have shed new light on their formation and dynamics. In a previous work, our lab demonstrated that insulator bodies form after dissociation of insulator proteins from chromatin and that insulator body formation can be induced under conditions of high osmotic pressure (Schoborg et al., 2013).

Immunostaining of S2 cells for γH2Av and Su(Hw) in isotonic cell culture media shows no visibly apparent pattern between the two proteins (Figure 4A). Linescans support this point, with little covariance seen between the signals (Figure 4A). Incubation of S2 cells in hypertonic media (containing an additional 250 mM NaCl) induced strong insulator body formation as previously reported (Schoborg et al., 2013). Immunostaining shows clear localization of γH2Av to insulator bodies (Figure 4B). Linescans drawn through the insulator bodies confirm this finding, with each signal following the same pattern (Figure 4B). The vast majority of observed insulator bodies formed during osmotic stress were positive for γH2Av (92.1%, Figure 4C). A number of smaller insulator bodies containing γH2Av also formed under isotonic conditions, implying that some amount of phosphorylated H2Av may be involved in insulator body function during normal cellular conditions. Alternatively, these cells may represent cells undergoing apoptosis (Schoborg et al., 2013). Quantitative colocalization analysis shows a clear and significant increase in colocalization between Su(Hw) and γH2Av after hypertonic stress, whereas there is no colocalization between the two signals in isotonic media (Figure 4D).

**Figure 4.**
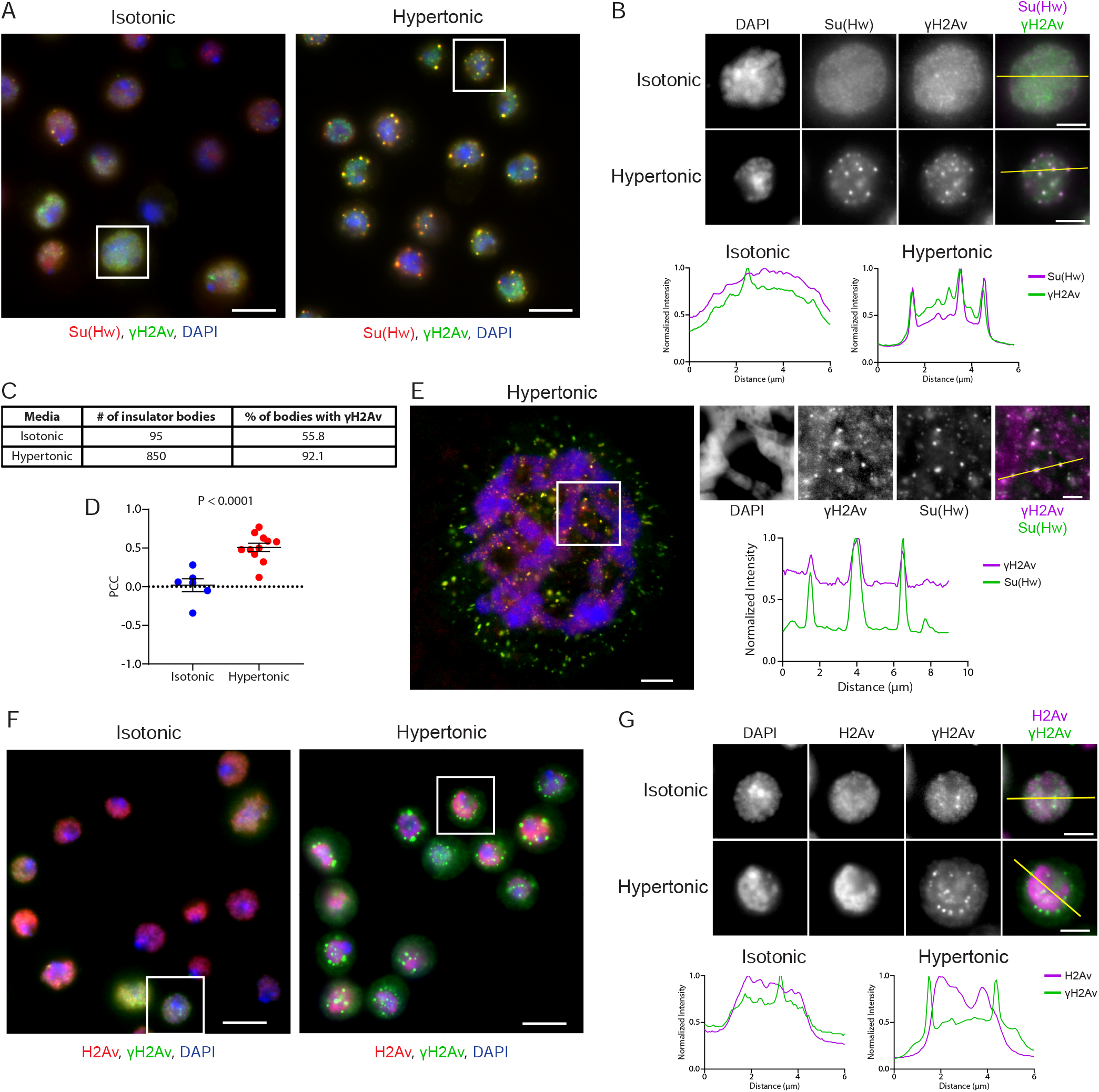
Phosphorylated H2Av is a component of insulator bodies. **A.** Immunostaining of *Drosophila* S2 cells in isotonic media (left) and hypertonic media (right). Insulator bodies formed during osmotic stress are labelled with Su(Hw) (red). Phosphorylated H2Av (green) colocalizes with Su(Hw) in insulator bodies. Scale bars are 5 μm and insets for analysis are delineated by white boxes. **B.** Magnified view of insets showing insulator body formation in hypertonic, but not isotonic, conditions. Insets are shown as RGB merge, with DAPI on the blue channel, γH2Av on the green channel, and Su(Hw) on the red channel. Red and green channels are shown independently in grey scale and merged as magenta and green. Beneath the insets are linescans corresponding to the yellow lines in the merged insets. Scale bars represent 2 μm. **C.** Table showing the ratio of insulator bodies with significant amounts of γH2Av. **D.** Pearson’s Correlation Coefficient (PCC) for γH2Av signal with Su(Hw) signal is plotted for isotonic versus hypertonic conditions, with each point representing a field of S2 cells. Error bars represent one standard error of the mean. The P-value was determined using an unpaired two-tailed Student’s T-test. **E.** Polytene chromosomes from wandering third instar larval salivary glands under hypertonic conditions. Colocalization between γH2Av and Su(Hw) is shown in the insets and linescan. The scale bar in the wide view represents 5 μm, the scale bar in the inset represents 2 μm. **F.** Immunostaining of *Drosophila* S2 cells in isotonic media (left) and hypertonic media (right). Phosphorylated H2Av (green) localizes to insulator bodies while unphosphorylated H2Av (red) does not. Scale bars are 5 μm and insets for analysis are delineated by white boxes. **G.** Magnified insets from F. Beneath the insets are linescans corresponding to the yellow lines in the merged insets. Scale bars represent 2 μm. Linescan intensities in B, E, and G were normalized by dividing each value by the maximum intensity recorded on each channel.

Insulator bodies have previously been shown to form in other cell types such as the polytene cells of the larval salivary gland (Schoborg et al., 2013). In order to determine whether or not phosphorylated H2Av was present in insulator bodies formed under osmotic stress conditions, larval salivary glands were treated in hypertonic solution and immunostained for Su(Hw) and γH2Av. Analysis of the micrographs reveals strong colocalization of the γH2Av antibody in the insulator bodies labelled by Su(Hw) in salivary gland cells (Figure 4E). There is no significant accumulation of unphosphorylated H2Av in these insulator bodies (Figure 4F and 4G), suggesting that phosphorylation of this histone variant is important for its localization to insulator bodies. The mechanism behind how insulators leave chromatin and form bodies is unknown, and the involvement of γH2Av adds an unexpected layer to this question. The presence of γH2Av in insulator bodies may also have implications in how cells respond to DNA damage.

### Phosphatase Inhibition Affects Interactions between γH2Av and *gypsy* Insulators

H2Av phosphorylation is tightly regulated by a network of kinases and phosphatases that integrate signals from various aspects of cellular activity including DNA damage repair (Sirbu and Cortez, 2013). Dephosphorylation of mammalian γH2AX is mitigated primarily through Protein Phosphatase 2A (PP2A), but also occurs through the activity of other phosphatases, including Protein Phosphatase 4 (PP4) (Nakada et al., 2008) and Wip1 phosphatase (Macurek et al., 2010). In order to determine how H2Av phosphorylation affects genome dynamics, we used okadaic acid (OA), a potent inhibitor of serine/threonine phosphatases PP1 and PP2A *in vitro* and *in vivo* (Bialojan and Takai, 1988, Haystead et al., 1989). As okadaic acid may affect other phosphatases at high concentrations (Honkanen and Golden, 2002), a concentration of 50 nM was chosen. Third-instar larval salivary glands were dissected then incubated in okadaic acid before fixation, squashing, and immunostaining. Examination of the polytene chromosomes showed no adverse effect of okadaic acid on the binding of insulator proteins or their colocalization with phosphorylated H2Av (Figure 5A, C, D). Quantification of the antibody signals indicates a significant increase in the amount of Su(Hw) signal in the presence of okadaic acid compared to untreated samples (Figure 5B). Notably, there is also an increase in the amount of phosphorylated H2Av bound to the polytene chromatin in the presence of okadaic acid compared to the untreated control (Figure 5B). This would seem to indicate that the okadaic acid is inhibiting PP2A from dephosphorylating γH2Av in chromatin, resulting in a significant accumulation of the modified histone variant. This is in conflict with findings from human cell culture, which were shown to increase cellular levels of H2AX phosphorylation in response to DNA damaging agents in the presence of 25 nM okadaic acid, but not in the presence of okadaic acid alone (Chowdhury et al., 2005). The difference may be a result either of the lower concentration of okadaic acid used in the previous study, or of the innate differences between the two model systems, i.e. the dual nature of *Drosophila* H2Av, which has the combined functions of mammalian H2AX and H2AZ (van Daal et al., 1988). The increased γH2Av signal in the presence of okadaic acid and in the absence of exogenous DNA damage may therefore support the H2AZ-like transcriptional regulation role in insulator function.

**Figure 5.**
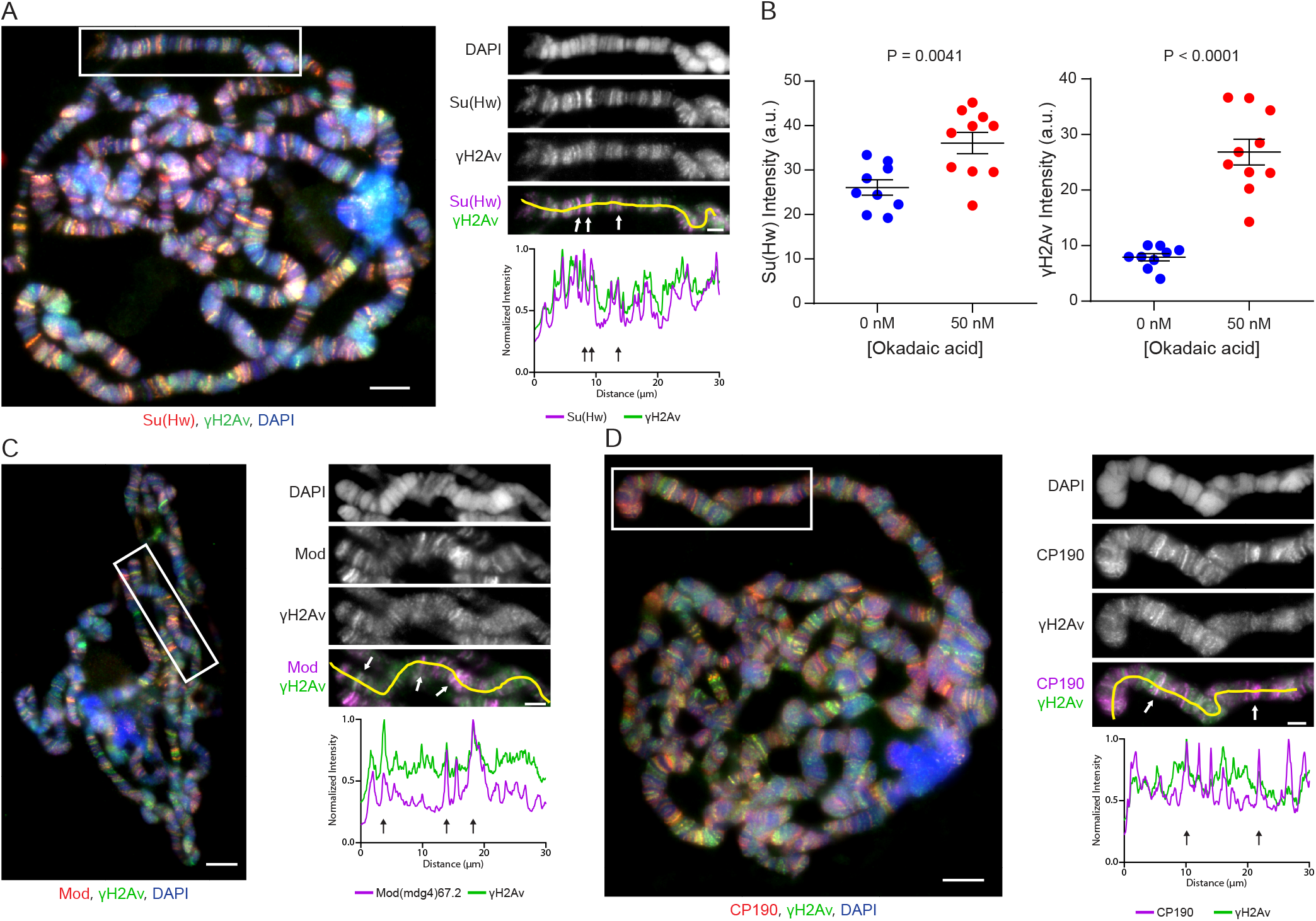
Treatment with okadaic acid increases the amount of phosphorylated H2Av on polytene chromosomes. Shown are co-immunostains of polytene chromosomes from salivary glands treated with okadaic acid. **A.** Immunostaining of γH2Av with Su(Hw). **B.** Quantification of the immunostaining data show in A. The intensities of Su(Hw) and γH2Av are shown under in the absence and presence of okadaic acid. Each point represents the polytene genome of an individual cell. Error bars represent one standard error of the mean. P-values were determined using unpaired two-tailed Student’s T-tests. **C.** Immunostaining of γH2Av with Mod(mdg4)67.2. **D.** Immunostaining of γH2Av with CP190. Immunofluorescent micrographs of polytene chromosome squashes are shown on the left. Magnified insets are shown to the right of each figure, corresponding to the white boxes in the figures on the left. Scale bars are 5 μm in the figures and 2 μm in the insets. Insets are shown as RGB merge, with DAPI on the blue channel, γH2Av on the green channel, and various insulator proteins on the red channel. Red and green channels are shown independently in grey scale and merged as magenta and green. Beneath the insets are linescans corresponding to the yellow lines in the merged insets. Linescan intensities in A, C, and D were normalized by dividing each value by the maximum intensity recorded on each channel.

Intriguingly, immunostaining of salivary glands from insulator mutants after incubation in okadaic acid revealed a significant rescue of both H2Av phosphorylation and its colocalization with components of the *gypsy* insulator complex. Close examination of the polytene chromosomes in the *su(Hw)^e04061^* mutant shows many sites of colocalization between γH2Av with both Mod(mdg4)67.2 and CP190 (Figure 6 A, B) which were lacking in the untreated mutant (Figure 3 A, B). This result suggests that in *su(Hw)* mutants, Mod(mdg4)67.2 interacts with CP190 and that this interaction is enhanced by γH2Av. Similar results were obtained when staining for Su(Hw) and CP190 in the *mod(mdg4)^u1^* mutant, with both proteins showing colocalization with phosphorylated H2Av after incubation with okadaic acid (Figure 6 C, D). As with the *su(Hw)^e04061^* mutant, this is in contrast to the untreated mutant samples in which γH2Av and *gypsy* insulator complexes do not colocalize. Staining of the *trans*-heterozygous *cp190^P11/H31-2^* mutant for Su(Hw) after okadaic acid treatment yielded the same response, with colocalization between Su(Hw) and γH2Av being rescued compared to the untreated mutant (Figure 6E). Colocalization analysis over the entire area of the polytene chromosomes showed strong correlation between the γH2Av signal and signals from insulator proteins in the okadaic acid-treated insulator mutants (Figure 6F). This is reflected in positive values for Pearson’s Correlation Coefficient (PCC), indicating significant correlations between γH2Av and *gypsy* insulator signals. This is in contrast to untreated mutant samples, which show a lack of correlation between γH2Av and *gypsy* insulator proteins (Figure 3F). These findings point to a model in which γH2Av acts as a component of *gypsy* insulators. Phosphorylation of H2Av may be required for *gypsy* insulator complex formation or stabilization, and may play an essential role in insulator functions.

**Figure 6.**
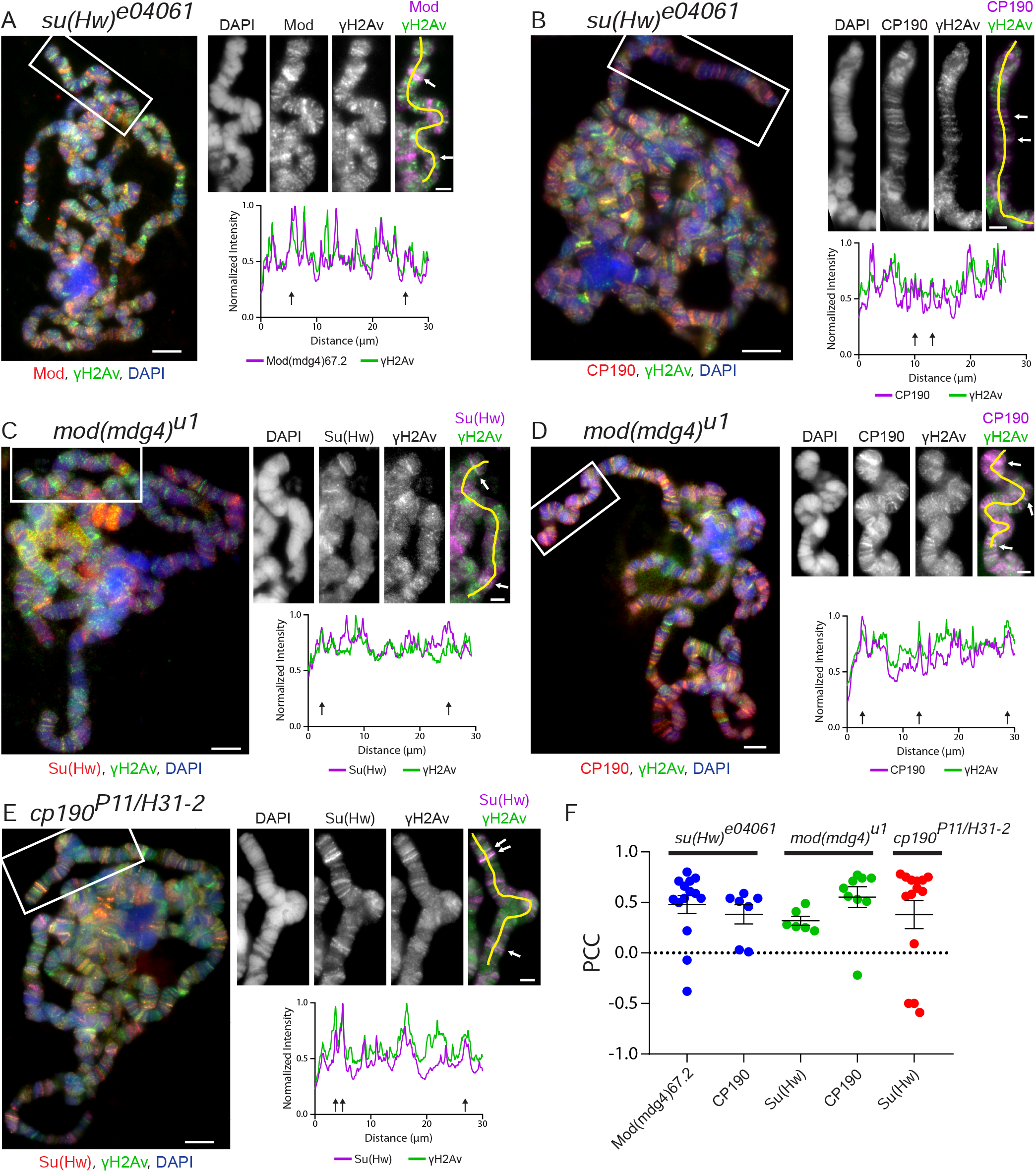
Phosphatase inhibition restores localization of γH2Av at insulator sites in insulator protein mutants. Shown are co-immunostains of polytene chromosomes from salivary glands treated with okadaic acid. **A.** Colocalization of γH2Av with Mod(mdg4)67.2 in *su(Hw)^e04061^*. **B.** Colocalization of γH2Av with CP190 in *su(Hw)^e04061^*. **C.** Colocalization of γH2Av with Su(Hw) in *mod(mdg4)^u1^*. **D.** Colocalization of γH2Av with CP190 in *mod(mdg4)^u1^*. **E.** Colocalization of γH2Av with Su(Hw) in *cp190^P11/H31-2^*. **F.** Pearson’s Correlation Coefficient (PCC) for γH2Av signal with each insulator protein signal is plotted, with each point representing the polytene genome of each cell. Error bars represent one standard error of the mean. PCC values are grouped by genotype (red = *su(Hw)^e04061^*, green = *mod(mdg4)^u1^*, blue = *cp190^P11/H31-2^*). Immunostaining results from polytene chromosome squashes are shown on the left in each panel. Magnified insets are shown to the right of each figure, corresponding to the white boxes in the figures on the left. Scale bars are 5 μm in the figures and 2 μm in the insets. Insets are shown as RGB merge, with DAPI on the blue channel, γH2Av on the green channel, and various insulator proteins on the red channel. Red and green channels are shown independently in grey scale and merged as magenta and green. Beneath the insets are linescans corresponding to the yellow lines in the merged insets. Linescan intensities were normalized by dividing each value by the maximum intensity recorded on each channel.

We next asked if this role of γH2Av in *gypsy* insulator dynamics is limited to chromatin-bound insulators or if it extends to other cellular functions of insulators. To this end, S2 cells were exposed to osmotic stress to induce insulator body formation with the goal of determining if H2Av phosphorylation is required for insulator body formation or recovery after stress (Figure 7A). As a control to ensure okadaic acid alone does not induce body formation, cells were incubated in isotonic media with okadaic acid. These cells showed similarly low numbers of insulator bodies per cell as cells in untreated isotonic media (Figure 7B). Osmotic stress was introduced by increasing the salt concentration (Schoborg et al., 2013), resulting in the formation of many insulator bodies (Figure 7A, C). No significant difference was seen in the ratio of cells that contained insulator bodies when comparing okadaic acid-treated and -untreated samples (Figure 7B). Of particular interest, however, is the finding that after the osmotic stress media is replaced with isotonic media, cells treated with okadaic acid recover significantly less than cells not exposed to okadaic acid, retaining a greater number of insulator bodies throughout recovery (Figure 7B, C). These results put into context our finding that γH2Av is present in insulator bodies (Figure 3) and imply that phosphorylated H2Av must be maintained in the insulator body as part of the normal osmotic response. Preventing dephosphorylation of γH2Av by phosphatase inhibition prevents resolution of insulator bodies during isotonic recovery, suggesting this is an essential part of the mechanism governing the cellular response to osmotic stress. It remains unclear whether H2Av is phosphorylated after recruitment into insulator bodies or if chromatin-bound H2Av is phosphorylated in response to osmotic stress before localizing to insulator bodies. A third possibility is that the γH2Av observed in insulator bodies originates from H2Av that is already phosphorylated through normal metabolic activity. Further experiments will be necessary to discern which of these models is correct.

**Figure 7.**
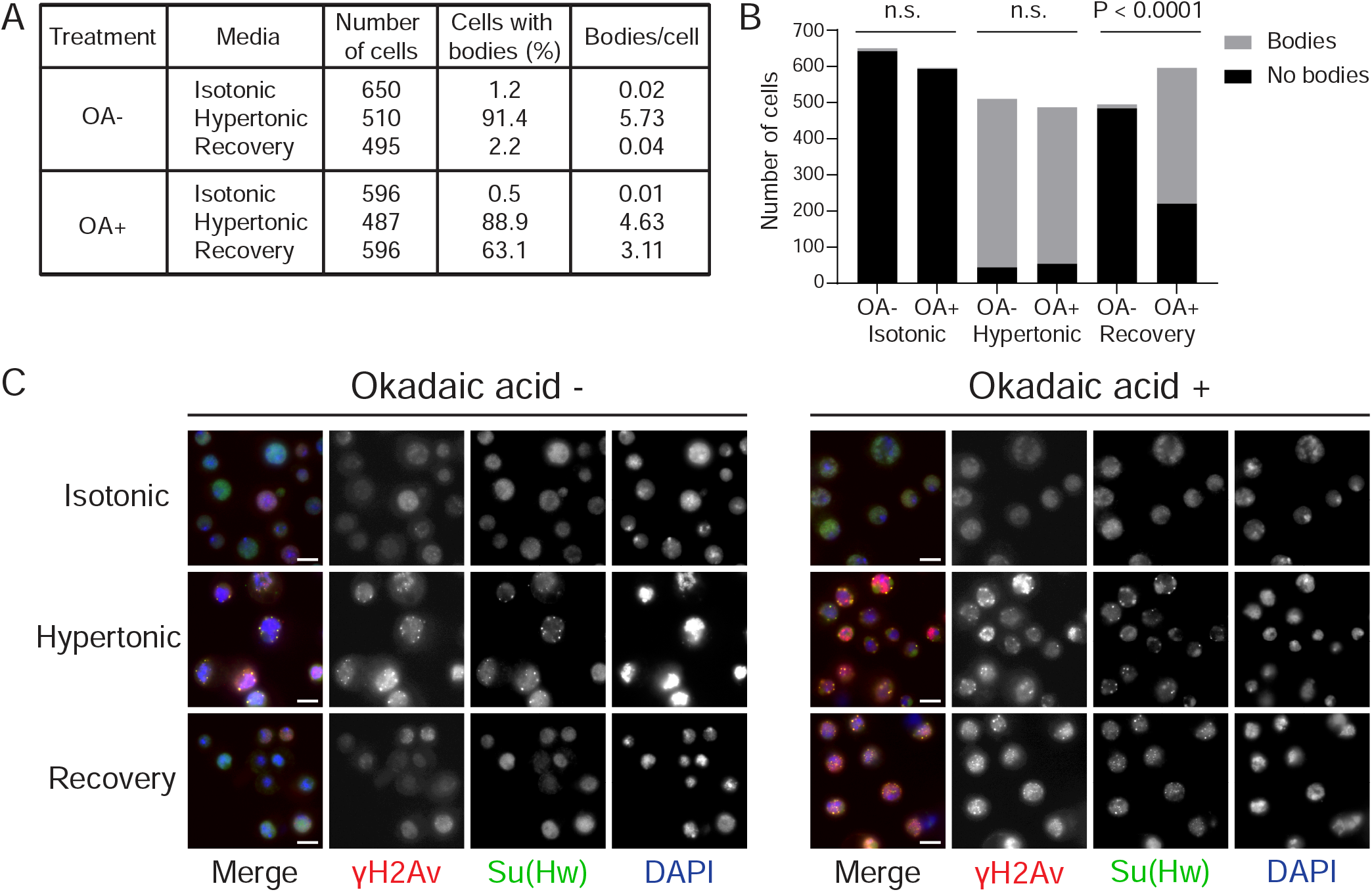
Phosphatase inhibition prevents recovery from insulator body formation atier osmotic stress. **A.** Tabulated results from OA (okadaic acid) treatment in osmotic stress and recovery. **B.** Bar graph showing the number of cells displaying bodies (grey) and those not displaying bodies (black). OA- and OA+ treatments are shown for each osmotic condition. P-values were determined using Fisher’s exact test. n. s. = not significant. **C.** Osmotic stress and recovery in the absence of okadaic acid (leti) in the presence of okadaic acid (right). Representative images are shown of S2 cells in isotonic media, in hypertonic stress media, and recovering in isotonic media. Merged images are shown on the leti, with each channel shown independently in greyscale. Scale bars represent 5 μm. Exposure times and contrast setings were kept constant between each sample.

### H2Av Is Phosphorylated in *gypsy* Insulator Sites

While a clear correlation is seen between H2Av phosphorylation and insulator binding proteins genome-wide, it is unclear if these interactions are a feature of insulators or sites where insulator proteins bind independently as transcription factors. For example, outside of the canonical *gypsy* insulator, Su(Hw) binds many sites in the genome alone or in conjunction with either CP190 or Mod(mdg4) (Kuhn-Parnell et al., 2008, Bushey et al., 2009, Soshnev et al., 2012, Soshnev et al., 2013). We therefore asked whether or not H2Av phosphorylation is required specifically at *gypsy* insulator sites by looking at two classical examples, *y^2^* and *ct^6^* (Figure 9A). The former is an allele of the *yellow* (*y*) gene in which a *gypsy* insulator site inserted into the region between enhancers for expression in the wing and body of the fly and the promoter, cutting off contact between the enhancers and promoter upstream of the *yellow* gene promoter and resulting in lowered expression of the *yellow* gene and ultimately lighter pigmentation of the adult fly (Harrison *et al.*, 1989, Geyer and Corces, 1992). The latter example is an allele of the *cut* (*ct*) gene in which a *gypsy* insulator between the wing margin enhancer and *cut* promoter prevents expression of the *cut* gene in the developing wing. This decreases *cut* expression in cells of the wing margin, leading to a jagged appearance of the wing margin (Jack *et al.*, 1991, Kim *et al.*, 1996). As it has previously been demonstrated that a functional gypsy insulator complex composed of Su(Hw), Mod(mdg4)67.2, and CP190 is required for proper function of the insulator (Georgiev and Kozycina, 1996, Cai and Levine, 1997, Gause *et al.*, 2001, Ghosh *et al.*, 2001, Mongelard *et al.*, 2002), these sites serve as known examples of genomic loci associated with binding of each of these components.

To determine if H2Av is phosphorylated at known *gypsy* sites, immunostaining of the X polytene chromosomes of larvae carrying the *y^2^* and *ct^6^* alleles were performed in wild-type and mutant backgrounds. Notably, a strong colocalization was observed for phosphorylated H2Av and each *gypsy* insulator component (Su(Hw), Mod(mdg4)67.2, and CP190) at both *y^2^* and *ct^6^* sites (Figure 8A). Strikingly, this colocalization is lost in *mod(mdg4)^u1^* mutants (Figure 8B). Su(Hw) is still recruited to *gypsy* insulator sites in *mod(mdg4)^u1^* as expected based on previous reports of *mod(mdg4)* mutants (Ghosh *et al.*, 2001, Melnikova, Kostyuchenko*, et al.*, 2017); more significant, however, is the observation that γH2Av is no longer observed colocalizing with Su(Hw) at *y^2^* or *ct^6^*. Likewise, H2Av phosphorylation is no longer observed at either *y^2^* or *ct^6^* in the *su(Hw)^e04061^* mutant background (Figure 8C). The lack of Mod(mdg4)67.2 at *y2* and *ct6* in the absence of Su(Hw) agrees with previous reports, which implicate Su(Hw) as necessary for recruitment of Mod(mdg4)67.2 to *gypsy* loci (Ghosh et al., 2001, Melnikova, Kostyuchenko, Molodina, et al., 2018), while the lack of γH2Av implies that either Su(Hw) is directly needed to maintain H2Av in a phosphorylated state or that a complete *gypsy* complex containing Mod(mdg4)67.2 is required. To expand on this question, polytene immunostaining from the *trans*-heterozygous *cp190^P11/H31-2^* mutant was performed (Figure 8D). This showed a decrease in the amount of Su(Hw) present at *y^2^* and *ct^6^*, consistent with a previous report that found reductions in both Su(Hw) and Mod(mdg4)67.2 in the polytene chromatin of *cp190* mutants, (Melnikova, Kostyuchenko, Molodina, et al., 2018). Similar to the mutations in *su(Hw)* and *mod(mdg4)*, mutation of *cp190* also significantly reduced levels of H2Av phosphorylation at the two *gypsy* loci examined. All together, these results indicate that reduction of any of the canonical *gypsy* insulator components is sufficient to disrupt H2Av phosphorylation at these sites. This would argue against this relationship being limited to Su(Hw)-only sites and supports the notion that H2Av phosphorylation is either promoted or stabilized by interactions with intact *gypsy* insulator complexes.

**Figure 8.**
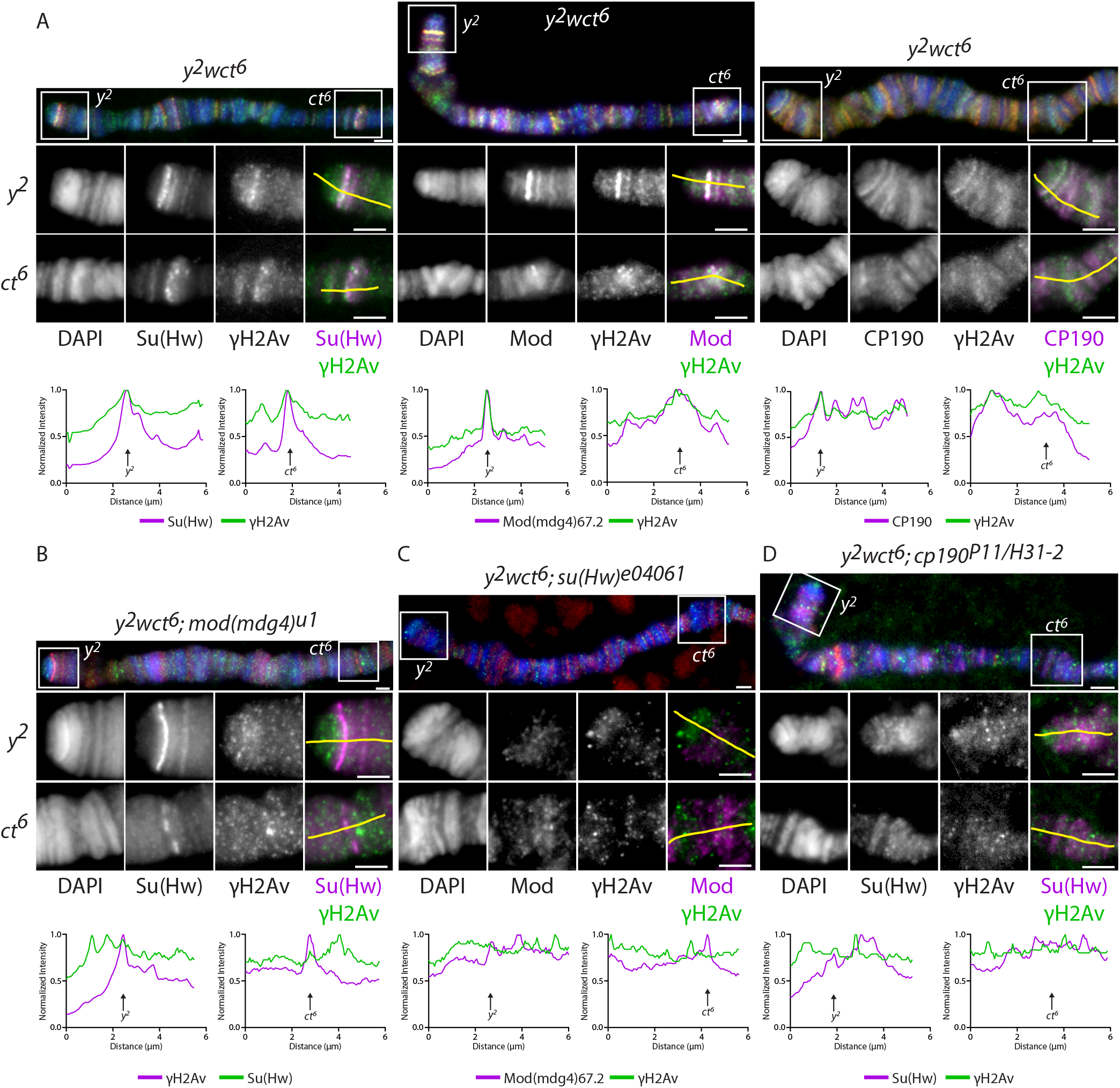
Phosphorylated H2Av is present at *gypsy* insulator sites. **A.** Colocalization between γH2Av and Su(Hw) (left), Mod(mdg4)67.2 (center), and CP190 (right) in a wild type *y^2^wct^6^* background. **B.** Colocalization between γH2Av and Su(Hw) in a *mod(mdg4)^u1^* homozygous background. **C.** Colocalization between γH2Av and Mod(mdg4)67.2 in a *su(Hw)^e04061^* homozygous background. **D.** Colocalization between γH2Av and Su(Hw) in a *cp190^P11/H31-2^ trans*-heterozygous background. Immunostaining results are shown of the X polytene chromosome from wandering third instar larvae in the *y^2^wct^6^* background. Each panel shows an X chromosome, with the *y^2^* and *ct^6^* sites labelled and delineated with white boxes. Beneath these are insets showing the *y^2^* and *ct^6^* sites in detail. Insets are shown as RGB merge, with DAPI on the blue channel, γH2Av on the green channel, and various insulator proteins on the red channel. Red and green channels are shown independently in grey scale and merged as magenta and green. Beneath the insets are linescans corresponding to the yellow lines in the merged insets. Arrows represent the approximate sites of each *gypsy* locus. Scale bars are 2 μm in the main figures and insets. Linescan intensities were normalized by dividing each value by the maximum intensity recorded on each channel.

### H2Av Contributes to *gypsy* Insulator Function

Our earlier results strongly suggest a correlation between γH2Av and *gypsy* insulator components, including colocalization on polytene chromatin (Figures 1 and 8) and in insulator bodies (Figure 4). Some evidence for a functional relationship was found in the dissolution of insulator bodies (Figure 7) but it remained unclear if γH2Av plays any role in canonical insulator functions such as enhancer-blocking. Based on the correlations described above between *gypsy* insulator proteins and γH2Av, we wondered whether or not mutation of *His2Av,* the sole H2A variant gene found in the *Drosophila* genome (van Daal et al., 1988), would affect *gypsy* insulator function. As the null mutation of *His2Av* is homozygous lethal at the third instar larval stage (van Daal and Elgin, 1992), we were precluded from observing how a complete lack of H2Av would affect the *yellow* and *cut* phenotypes seen in adults. We therefore set up a series of crosses to examine how adult animals that carried heterozygous mutations for both *His2Av* and *su(Hw)*.

The effects of these mutants on the *ct^6^* allele were measured by examining the wing margins. Wing margins in *Drosophila* contain mechanosensory bristles (Hartenstein and Posakony, 1989) that require expression of the *cut* gene for proper formation (Jack, 1985). Lack of *cut* expression results in decreased specification and differentiation of mechanosensory bristles and increased rates of cell death (Jack et al., 1991, Liu et al., 1991). The *ct^6^* phenotype is characterized by wings having an incomplete margin due to such defects, resulting in the nominal “cuts” in the wing which can be rescued to near wild type by mutations in *gypsy* insulator genes (Georgiev and Kozycina, 1996, Kim et al., 1996). Wing width was measured as a proxy for *cut* gene activity, as wings with the *ct^6^* phenotype are less wide due to the decreased cell proliferation associated with this phenotype. Width of the wing is calculated as the area divided by the feret diameter (a measure of length). Notably, flies that contained heterozygous mutations for both *His2Av^810^* and *su(Hw)^e04061^* displayed wider wings, while both single heterozygotes showed varying degrees of cuts in the margin (Figure 9B). To verify this finding, the null *su(Hw)^V^* allele was next tested in conjunction with *His2Av^810^*. Consistent with the *His2Av^810^/su(Hw)^e04061^* genotype, *His2Av^810^/su(Hw)^V^* double heterozygotes had significantly wider wings than either single heterozygote (Figure 9B). This increased suppression of *gypsy* insulator phenotypes in *su(Hw)* heterozygous backgrounds implies that H2Av is involved in *gypsy* insulator function.

**Figure 9.**
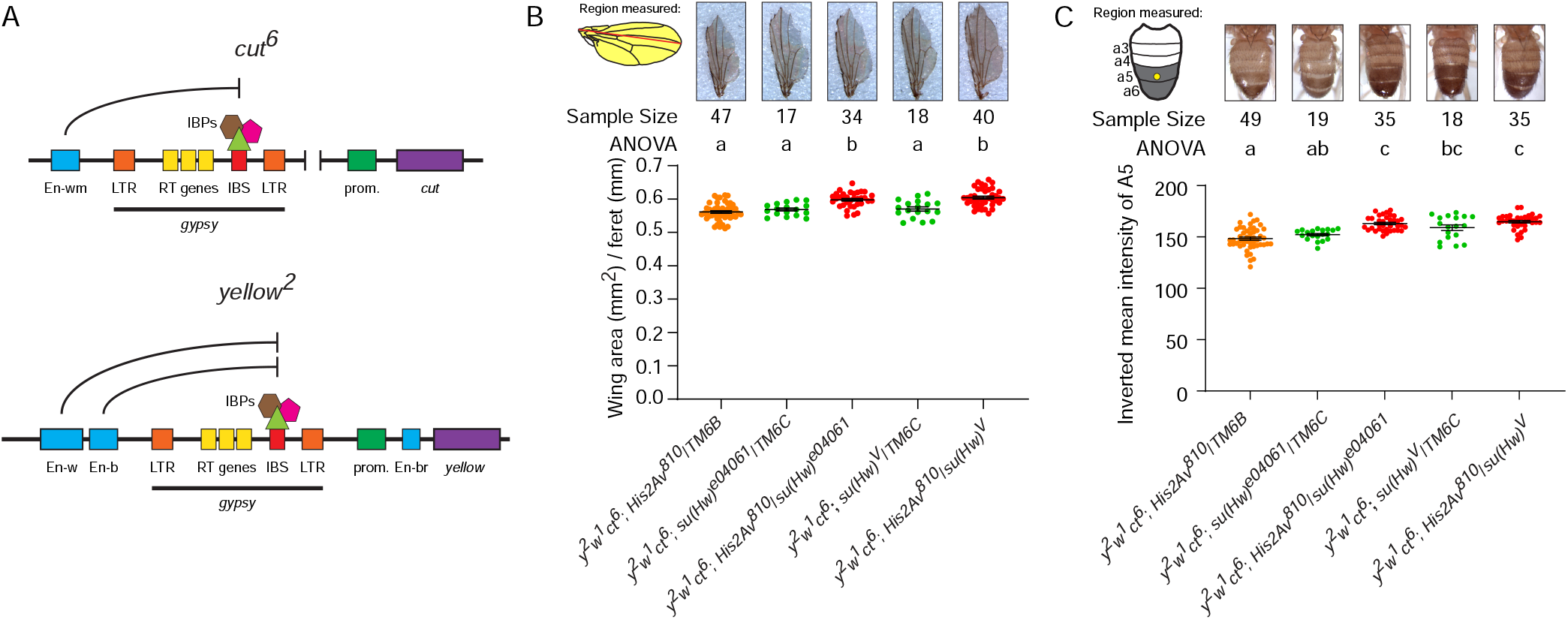
H2Av contributes to *gypsy* insulator function. **A.** Illustration of the upstream regulatory elements found in the *ct^6^* and *y^2^* alleles. Gene coding sequences (purple) are regulated by coordinated contacts between enhancers (blue) and promoters (green). In each case, the *gypsy* retrotransposon (denoted with the thick black bar) has been inserted between a promoter and at least one enhancer. LTR = long terminal repeats, RT genes = retrotransposon genes (*gag*, *pol*, and *env*), IBS = insulator binding site, IBPs = insulator binding proteins (Su(Hw), Mod(mdg4)67.2, and CP190). When the IBS is bound by a complete insulator complex, it interrupts communication between the promoter and distal enhancers. In *cut^6^* this prevents expression of *cut* by the wing margin enhancer (En-wm), while in *yellow^2^* the insulator prevents expression of *yellow* directed by the wing enhancer (En-w) and the body enhancer (En-b) but not the bristle enhancer (En-br) between the *yellow* promoter and transcription start site. Genomic distances are not drawn to scale. **B.** The average length of wings along the feret axis was used as a metric for the *cut* phenotype. Wing area in mm^2^ (yellow shaded area) was divided by feret diameter (red line). Examples of wings are displayed above their respective genotypes. **C.** *y^2^* phenotype scoring in the male abdomen. The illustration on the leti shows the male abdomen with abdominal segments 3-6 labelled. A circular ROI (yellow circle) was sampled from the a5 tergite of each male to determine the degree of pigmentation. The graph depicts the mean pixel intensity from individual flies by genotype. Example pictures of abdomens are displayed above their respective genotypes. In B and C, green dots represent *su(Hw)* heterozygotes arising from each cross, orange dots represent *His2Av^810^* heterozygotes, and red dots indicate flies doubly heterozygous for *His2Av^810^* and *su(Hw)*. Above each data set is the sample size and single letier statistical codes from an ANOVA performed with Dunneti’s T3 multiple comparisons, with significant differences at P ≤ 0.05.

In order to determine if this requirement for H2Av in *gypsy* insulator function is specific to the *cut* gene or if it is a general mechanism for *gypsy* insulators, we tested the effect of *His2Av* mutation on the *y^2^* phenotype. Effects of these mutations on the *y^2^* allele were tested by measuring the degree of pigmentation in the wings and in the darkened A5 tergite found in male flies. Heterozygous *HisAv^810^* or *su(Hw) ^e04061^* mutations by themselves have little effect on expression of *yellow* in the abdomen (Figure 9C). In contrast to this, abdomens in the double heterozygous *His2Av^810^/su(Hw)^e04061^* mutants were significantly darker than the single heterozygous mutants (Figure 9C). This implies a reduction in the enhancer-blocking capacity of the *gypsy* insertion upstream of *yellow* and suggests a functional role for H2Av in *gypsy* insulator function. In order to exclude the possibility that second-site mutations in the *su(Hw)^e04061^* background were influencing this rescue, we crossed the null *su(Hw)^V^* allele with the *His2Av^810^* mutant. Flies doubly heterozygous for *su(Hw)^V^* and *His2Av^810^* showed a statistically significant increase in pigmentation compared to the *His2Av^810^* heterozygote, but not the single *su(Hw)^V^* mutant (Figure 9C). This discrepancy between *su(Hw)^e04061^* and *su(Hw)^V^* may be due to mutation of the neighboring *RpII15* gene (an RNA Pol II subunit) in the *su(Hw)^V^* chromosome (Harrison et al., 1992), which may have an epistatic effect on this phenotype. Taken together, these findings represent the first evidence for the requirement of a histone protein in *Drosophila* insulator function and may hint at a mechanism for how insulators work. There is a correlation between the lack of a balancer chromosome and the rescues seen in this assay. The lack of a balancer is unlikely to be a causative factor in the rescues, as the TM6B and TM6C balancers present in the single heterozygotes carry the *ebony^1^* marker, which significantly increases the degree of pigmentation (Wittkopp *et al.*, 2002), and the single heterozygotes are lighter than the trans-heterozygotes.

### γH2Av colocalizes with Su(Hw) at TAD boundaries

Next, given the previous observations indicating that γH2Av and Su(Hw) sites colocalize in polytene chromosomes, we asked whether the colocalization of both proteins is also supported at the molecular level by Chromatin Immunoprecipitation (ChIP) experiments. To answer this question, we used publicly available ChIP data for γH2Av and Su(Hw) from Kc167 cells (SRX1299942 SRA) (Li et al., 2016). We used SeqMiner to compare the two ChIP-seq datasets by heatmap analysis. The peak summits for γH2Av were taken as the reference coordinate for a heatmap and profile analysis comparing Su(Hw) and γH2Av distributions (Figure 10A). The profile of mean read densities for both proteins also shows a significant overlap between Su(Hw) and γH2Av peaks at γH2Av peaks flanked by 10,000 bp (Figure 10B). These results support our observation that the genomic distribution of Su(Hw) and γH2Av overlap in polytene chromosomes.

**Figure 10:**
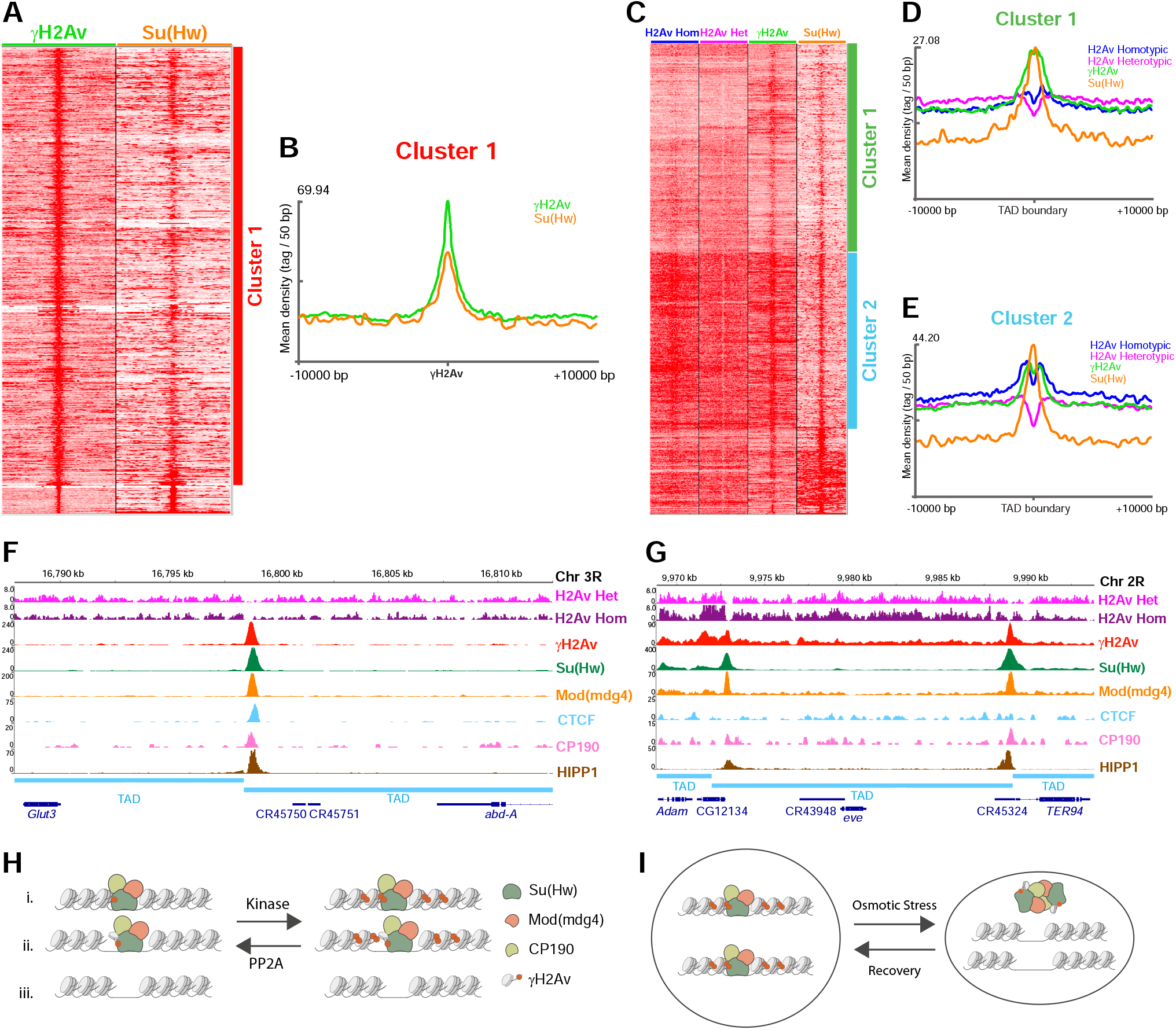
Phosphorylated H2Av associates with TAD boundaries. **A.** Heatmap comparing the intensity distributions of γH2Av and Su(Hw) using γH2Av peaks as reference. **B.** Mean read density profiles of γH2Av and Su(Hw) centered at γH2Av peaks. **C.** Heatmap comparing the intensity distributions of H2Av nucleosomes, γH2Av, and Su(Hw) using TAD boundaries as a reference. **D.** and **E.** Mean read density profiles of two major clusters from heatmap in C, centered at TAD boundaries. **F.** Peak profile of insulator proteins, including γH2Av and the H2Av nucleosome distribution at the leti boundary of the *Abd-A* TAD. **G.** Peak profile of insulator proteins, including γH2Av and the H2Av nucleosome distribution at the Homie insulator flanking the pair rule gene *eve*. **H.** This model shows the interaction between *gypsy* insulator components and H2Av phosphorylation. We propose that binding of the Su(Hw) insulator complex promotes phosphorylation of neighboring H2Av. This stabilizes binding of the insulator complex and promotes spread of the H2Av phosphorylation mark through neighboring nucleosomes. (i.) Case in which Su(Hw) complexes interact with nucleosomal γH2Av. (ii.) Case in which Su(Hw) complexes interact with non-nucleosomal γH2Av. (iii.) Case in which Su(Hw) is absent and phosphorylation of H2Av is significantly reduced. **I.** During osmotic stress, insulator complexes leave chromatin and form insulator bodies. These bodies contain γH2Av that must be dephosphorylated during recovery so that insulator proteins can return to their genomic binding sites.

Because insulator proteins are frequently associated with the boundaries of Topologically Associating Domains (TADs), we next asked whether γH2Av is also enriched at TAD boundaries. To address this question, we used publicly available data and obtained a map of all *Drosophila* TADs as determined by Hi-C using *Drosophila* Kc167 cells (Ramirez et al., 2018). We used this data to generate a BED file consisting of 1,000 bp DNA fragments containing all TAD boundaries in the *Drosophila* genome flanked by 500 bp at both sites of the boundary. The 1 kb TAD boundary BED file was used as a reference coordinate for a heatmap comparing the density distribution of nucleosomes containing the histone variant H2Av, either homotypic (H2Av Hom) or heterotypic (H2Av Het), as well as the densities of yH2Av and Su(Hw) in the *Drosophila* genome (Figure 10C). The distribution of H2Av nucleosomes was obtained from a previous study (Weber et al., 2010). The heatmap analysis shows there is a strong association between the density distribution of H2Av nucleosomes and TAD boundaries (Figure 10C). Generally, we found homotypic H2Av nucleosomes are enriched at the boundaries, whereas heterotypic nucleosomes have a significant density drop at TAD boundaries.

Interestingly, enrichment in γH2Av and Su(Hw) is also observed at the TAD boundaries (Figure 10C). Signal enrichment can be grouped in two major clusters, where the most significant difference is the relative enrichment of Su(Hw) and γH2Av at the boundaries (Figure 10D, E). In cluster 2, heterotypic H2Av and γH2Av have similar enrichment levels through the DNA flanking the boundary. At the center of the boundary, however, is the homotypic instead of the heterotypic H2Av that has an enrichment similar to that of γH2Av, with both intensity profiles (H2Av Hom and γH2Av) significantly more elevated than in the flanking DNA (Figure 10E). Interestingly, Cluster 1 has the opposite pattern. In cluster 1, homotypic H2Av and γH2Av have similar enrichment densities through the DNA flanking the boundary and remain elevated at the boundary center. However, γH2Av and Su(Hw) intensities are similar and higher than that of the homotypic and heterotypic H2Av nucleosomes (Figure 10D).

The functional significance of the association of γH2Av with TAD boundaries is intriguing. Interestingly, we found that the association of γH2Av with TAD boundaries is very similar to that of Su(Hw) and other insulator proteins in that is found as an enriched peak. One such example of this is a TAD boundary that flanks a TAD containing the homeotic gene *Abdominal A* (*Abd-A*). Like other boundaries associated with the homeobox gene cluster (Postika *et al.*, 2018, Ozdemir and Gambetta, 2019), this boundary is enriched in the insulator proteins Su(Hw), Modifier of Mdg4, CP190, CTCF, and HIPP1. Here, we show this boundary is equally enriched in γH2Av (Figure 10F). Moreover, when observing the distribution of H2Av nucleosomes at this site, it appears that the γH2Av peak does not colocalize with a nucleosome, suggesting the possibility that γH2Av may be non-nucleosomal. In another example, two boundaries flank a TAD that contains the developmentally regulated pair rule gene *eve* (Figure 10G). Each boundary is enriched in γH2Av; however, in the left boundary the γH2Av peak overlaps at least partially with the nucleosomal H2Av, whereas in the right boundary the γH2Av peak seems to be non-nucleosomal as well. This particular boundary corresponds to the well characterized Homie insulator (Fujioka et al., 2013). These findings show that the distribution of γH2Av in the *Drosophila* genome display all the properties of insulator proteins, suggesting that γH2Av may be required for insulator function.

## Discussion

Our results have demonstrated a clear relationship between the phosphorylated histone variant γH2Av and chromatin insulator proteins. γH2Av colocalizes with Su(Hw) insulator complexes throughout the genome, and both are enriched at TAD boundaries. Disruption of insulator complex formation prevents stable phosphorylation of H2Av that can be overcome by phosphatase inhibition. Importantly, we provide evidence that this histone variant is involved in insulator activity, as dephosphorylation of γH2Av is necessary for dissolution of insulator bodies and reducing the genetic dose of H2Av in a sensitized *su(Hw)* heterozygous mutant background partially rescues *gypsy* insulator phenotypes.

Chromatin insulator proteins were initially characterized by their enhancer-blocking properties and their ability to prevent the spread of heterochromatin, and more recently by their role in large-scale genome organization (Harrison *et al.*, 1989, Geyer and Corces, 1992, Bell *et al.*, 1999, Bushey *et al.*, 2009). In addition to these canonical properties, however, our lab has uncovered roles of insulators in other aspects of cell metabolism including the osmotic stress response (Schoborg *et al.*, 2013) and genome stability (Hsu *et al.*, 2019). It is now established that mutation of the only insulator protein found in humans, CTCF, predisposes cells to cancer formation (Docquier *et al.*, 2005, Kemp *et al.*, 2014, Guo *et al.*, 2018) through increased rates of unrepaired DNA damage. Despite this knowledge, the mechanisms linking insulator activity to the maintenance of genome stability remain enigmatic. Here, we report a functional relationship between a *Drosophila* insulator and H2Av, the sole histone H2A variant in fruit flies. H2Av in *Drosophila* performs functions associated with both the mammalian histone variants H2AX and H2AZ (Baldi and Becker, 2013), and misregulation of H2Av in *Drosophila* is associated with severe developmental phenotypes including nuclear falling in embryos (Li *et al.*, 2014), the formation of necrotic tumors in the larval lymph gland (Grigorian *et al.*, 2017) and an inability to maintain stem cell populations in adult tissues (Morillo Prado *et al.*, 2013). Like H2AX, H2Av is phosphorylated in response to DNA double strand breaks (DSBs) and serves as a chromosomal mark to recruit DNA repair proteins (Madigan *et al.*, 2002, Joyce *et al.*, 2011).

Our initial experiments in *Drosophila* polytene chromosomes revealed a striking correlation between the binding sites of phosphorylated H2Av and insulator proteins at *gypsy* insulator sites (Figure 8) and at Su(Hw) sites elsewhere in the genome (Figures 1, 10). Importantly, this colocalization depends on having a complete and stable Su(Hw) insulator complex, as mutating any of the three core insulator components reduces the coincidence of γH2Av signals with the remaining insulator proteins (Figure 3). Some insight into the mechanism behind this phenomenon comes from experimental inhibition of PP2A, the phosphatase responsible for dephosphorylating γH2Av after resolution of double strand breaks (Merigliano *et al.*, 2017). We found that the amount of γH2Av in undamaged polytene chromosomes increases after PP2A inhibition. This was as expected due to the known role of PP2A in regulating H2Av phosphorylation. Surprisingly, the amount of Su(Hw) bound to the polytene chromosomes also significantly rose when tissues were treated with the PP2A inhibitor (Figure 5). This seems to imply that *gypsy* insulator complex formation or stability is driven at least in part by the phosphorylation state of H2Av, although we cannot rule out the potential impact that other PP2A substrates may have on insulator complexes. Indeed, we show that inhibition of PP2A rescues *gypsy* insulator complex formation in tissues missing one of the three core insulator binding proteins (Figure 6). This supports the notion that Su(Hw) insulators are stabilized by the presence of phosphorylated H2Av.

To further explore the relationship between insulator binding proteins and γH2Av, we performed immunostains on polytene chromosomes in the background of mutations in genes encoding for insulator proteins. We observed significant reductions of γH2Av signal in *su(Hw)^e04061^* and *cp190^P11/H31-2^* (Figure 2), suggesting a relationship between γH2Av and *gypsy* insulator components in terms of recruitment to the chromatin. Notably, the reduction of γH2Av in *su(Hw)^e04061^* was more extreme than in other mutants. From this we postulate that the interaction between γH2Av and Su(Hw) insulator sites may be largely mediated through interactions with Su(Hw) itself, although interactions with Mod(mdg4)67.2 and CP190 may also contribute to the overall stability of the complex. A large fraction of Su(Hw) binding sites do not include Mod(mdg4)67.2 or CP190 based on chromatin immunoprecipitation data (Negre *et al.*, 2010). It will thus be informative for future experiments to determine whether or not the presence of H2Av or γH2Av is required for insulator activity or transcriptional regulation at other loci bound by Su(Hw).

Our results to this point showed numerous correlations between *gypsy* insulators and γH2Av in chromatin, but it remained unclear if there was a functional relationship. Therefore, our final round of experiments sought to determine if γH2Av influences insulator function. These experiments are limited by the lethality of adults H2Av homozygous mutant, but one key finding is the partial rescue of the *ct^6^* phenotype in *su(Hw)^e04061^*/*His2Av^810^* double heterozygotes (Figure 9). Significant increases in pigmentation in the abdomens of male flies were also found in this genotype in the *y^2^* background, and the *ct^6^* phenotype rescue was replicated in the *su(Hw)^V^*/*His2Av^810^* background (Figure 9). This demonstrates that H2Av influences the activity of insulator complexes in multiple tissues and therefore could be considered an insulator protein itself.

Our findings point to a model in which *gypsy* insulator components and γH2Av stabilize each other in the chromatin (Figure 10H). Based on our results and the fact that Su(Hw) makes direct interactions with DNA while Mod(mdg4)67.2 and CP190 are recruited by interactions with Su(Hw) (Harrison *et al.*, 1993, Gdula *et al.*, 1996, Ghosh *et al.*, 2001, Pai *et al.*, 2004, Melnikova, Kostyuchenko*, et al.*, 2017), we propose that Su(Hw) is the main site of the interaction with γH2Av and that Mod(mdg4)67.2, and potentially CP190, stabilize this interaction. Further biochemical analysis will be required to determine if γH2Av is in direct physical contact with Su(Hw) and, if so, which domains are the contact points between these proteins. Another possibility is an indirect association between γH2Av and Su(Hw), possibly via another yet unknown protein. Interactions with RNA may be another mechanism by which these proteins associate based on the recent finding that *Shep* RNA is required for *gypsy* insulator function (Chen *et al.*, 2019). Direct binding with Mod(mdg4)67.2 or CP190 cannot be ruled out either, though such interactions are likely to have less effect on γH2Av recruitment than that with Su(Hw), based on results from the *ct^6^* and *y^2^* phenotypic rescue experiment (Figure 9).

Our model also proposes that γH2Av localizes in insulator bodies that form as a result of osmotic stress (Figure 10I). The mechanism remains unknown with possibilities including disassembly of the complex and reassociation as nucleoplasmic bodies or the entire complex translocating from chromatin into insulator bodies. It is not yet clear, however, if the γH2Av found in insulator bodies comes from the same population as found in chromatin or if these proteins are recruited from the non-chromatin bound nucleoplasmic H2Av population.

An interaction between Su(Hw) and nucleosomes was recently reported in which *gypsy* insulators serve to position nucleosomes and establish regular nucleosome phasing (Baldi *et al.*, 2018). Our observations may relate to this finding if such Su(Hw)-based nucleosome positioning interactions depend on the presence of γH2Av. Another possibility is a potential relationship between contacts with *gyspy* insulator proteins and whether neighboring nucleosomes are heterotypic or homotypic for this histone (Weber *et al.*, 2010). Our ChIP-seq analysis shows enrichment for homotypic H2Av along with Su(Hw) and other insulator proteins at TAD borders (Figure 10). This raises the question of whether *gypsy* insulator proteins affect the composition of nearby nucleosomes, or conversely, if these insulator proteins are preferentially recruited to homotypic nucleosomes. It is also possible that H2Av is acting as an insulator protein outside of a nucleosome, as ChIP-seq peaks enriched for γH2Av and insulator proteins do not necessarily align to nucleosomes (Figure 10). While the mechanistic relationship between these groups of proteins remains unknown, our data add to a growing consensus that interactions between histone proteins and insulator proteins are required for cellular homeostasis and maintenance of genome integrity.

Recent results from mammalian cell culture studies highlight an interaction between H2AZ, the histone variant associated with transcriptional regulation (Giaimo *et al.*, 2019), and CTCF, the sole insulator protein in mammals (Wen *et al.*, 2020). As *Drosophila* H2Av serves orthologous functions to both mammalian H2AZ and H2AX, our findings regarding insulators and the early DSB marker γH2Av may be relevant to human health as CTCF is frequently mutated in various cancers (Kemp *et al.*, 2014, Canela *et al.*, 2017, Guo *et al.*, 2018). Mutation of CTCF in *Drosophila* is associated with instability in ribosomal DNA found in the nucleolus (Guerrero and Maggert, 2011). If the relationship between insulator proteins and the DNA damage repair pathway is conserved from flies to humans, testing for interdependence of these components may provide clinically useful information. Indeed, it was recently described that CTCF sets the boundaries for phosphorylated H2AX spreading in human cell culture (Natale *et al.*, 2017) and that CTCF and γH2AX are both recruited to sites of DSBs in mouse embryonic fibroblasts (MEFs) (Lang *et al.*, 2017). Unlike our findings with Su(Hw), CTCF depletion in MEFs increases H2AX phosphorylation but, similar to Su(Hw) depletion in flies, induces genome instability (Lang *et al.*, 2017), possibly indicating a different mechanism linking CTCF to H2AX phosphorylation. The link between DNA damage repair and insulator proteins is further corroborated by the recent finding that TAD boundary strength and CTCF insulation strength both increase in an ATM-dependent manner in human cell culture after X-ray-induced DNA damage (Sanders *et al.*, 2020).

Other questions generated from our results here will need to be addressed in future experiments. One example is determining how H2Av or its phosphorylated form affect other phenotypes associated with lack of *su(Hw)* expression. Importantly, *su(Hw)* has been shown to regulate genome stability (Lankenau *et al.*, 2000) and we recently described the presence of chromosomal aberrations in actively dividing neuroblasts of *su(Hw)*-deficient larvae (Hsu *et al.*, 2020). How these aberrations arise is still unclear, however, our finding here that *su(Hw)* homozygotes show significantly less phosphorylated H2Av may shed light on a mechanism. Cells that are unable to phosphorylate H2Av in response to DNA damage may be unable to recruit essential DNA damage repair proteins to the site of DSBs. If left alone, these unrepaired DSBs become obvious candidates for the source of the chromosomal aberrations seen in larval neuroblasts. Further studies will address this question, along with examining the role of Su(Hw) in repair of induced DNA damage, as well as the role of γH2Av in boundary function and genome organization. Our results so far provide a foundation for understanding how the interplay between chromatin insulators and histones influences gene regulation and genome stability.

## Acknowledgements

We would like to thank former members of the Labrador lab, including Dr. Emily Stow, for collaboration and discussion. We thank Dr. Rachel Patton McCord for critical review of the manuscript. Stocks obtained from the Bloomington *Drosophila* Stock Center (NIH P40OD018537) were used in this study. S2 cell culture was obtained from the *Drosophila* Genomics Resource Center (NIH 2P40OD010949).

## Author contributions

Conceptualization: J.R.S., R.A., M.L.; Data curation: J.R.S., R.A.; Formal analysis: J.R.S., B.A., M.L.; Funding acquisition: M.L.; Investigation: J.R.S., R.A., B.A., S.Z., J.K., M.L.; Methodology: J.R.S., R.A., M.L.; Project administration: M.L.; Resources: J.R.S., M.L.; Software: J.R.S.; Supervision: M.L.; Validation: J.R.S.; Visualization: J.R.S., B.A., M.L.; Writing – original draft: J.R.S., B.A., M.L.; Writing – review & editing: J.R.S., M.L.

